# The Spatial Atlas of Human Anatomy (SAHA): A Multimodal Subcellular-Resolution Reference Across Human Organs

**DOI:** 10.1101/2025.06.16.658716

**Authors:** Jiwoon Park, Roberto De Gregorio, Erika Hissong, Elif Ozcelik, Nicholas Bartelo, Felipe Segato Dezem, Luke Zhang, Maycon Marção, Hannah Chasteen, Yimin Zheng, Ernesto Abila, Junbum Kim, Theodore Nelson, Jacqueline Proszynski, Akua A. Agyemang, Mohith Reddy Arikatla, Arjumand Wani, Yutian Liu, Evelyn Metzger, Stefan Rogers, Prajan Divakar, Parambir S. Dulai, Jason Reeves, Yan Liang, Liuliu Pan, Sayani Bhattacharjee, Michael Patrick, Kimberly Young, Ashley Heck, Mithra Korukonda, Dan McGuire, Lidan Wu, Aster Wardhani, Joseph Beechem, George Church, Steven M Lipkin, Aaron Berliner, Sanjay Patel, Fabio Socciarelli, Jan Krumsiek, Rohit Chandwani, Sebastien Monette, Brian Robinson, Massimo Loda, Olivier Elemento, Luciano Martelotto, Jasmine Plummer, André F. Rendeiro, Alicia Alonso, Robert E. Schwartz, Shauna Lee Houlihan, Christopher E. Mason

**Affiliations:** Weill Cornell Medicine, New York, NY, USA; Harvard Medical School, Boston, MA, USA; Caryl and Israel Englander Institute for Precision Medicine, Weill Medical College of Cornell University, New York, USA; Department of Pathology and Laboratory Medicine, Weill Medical College of Cornell University, New York, NY, USA; Center for Spatial Omics, St. Jude Children’s Research Hospital, Memphis, TN, USA; Department of Developmental Neurobiology, St. Jude Children’s Research Hospital, Memphis, TN, USA; CeMM Research Center for Molecular Medicine of the Austrian Academy of Sciences, Vienna, Austria; Ludwig Boltzmann Institute for Network Medicine at the University of Vienna, Austria; Bruker Spatial Biology, Seattle, WA, USA; Department of Medicine, Division of Gastroenterology and Hepatology, Northwestern University Feinberg School of Medicine; Center for Human Immunobiology, Northwestern University Feinberg School of Medicine; Memorial Sloan Kettering Cancer Center, New York, NY, USA; University of Adelaide, Adelaide, AustCralia; Comprehensive Cancer Center, St. Jude Children’s Research Hospital, Memphis, TN, USA; Department of Cellular & Molecular Biology, St. Jude Children’s Research Hospital, Memphis, TN, USA

## Abstract

The Spatial Atlas of Human Anatomy (SAHA) represents the first multimodal, subcellular-resolution reference of healthy adult human tissues across multiple organ systems. Integrating spatial transcriptomics, proteomics, and histological features across over 15 million cells from more than 100 donors, SAHA maps conserved and organ-specific cellular niches in gastrointestinal and immune tissues. High-resolution profiling using CosMx SMI, 10x Xenium, RNAscope, GeoMx DSP, and single-nucleus RNA-seq reveals spatially organized cell states, rare adaptive immune populations, and tissue-specific cell-cell interactions and ligand-receptor pairs. Comparative analyses with colorectal cancer and inflammatory bowel disease demonstrate the power of SAHA to detect disease-associated spatial disruptions, including crypt dedifferentiation, perineural invasion, and therapy-resistant immune remodeling. All data are openly accessible through a FAIR-compliant interactive portal to support exploration, benchmarking, and machine learning model training. Through SAHA, we provide a foundational framework for spatial diagnostics and next-generation precision medicine grounded in a comprehensive human tissue atlas, enabling the development of context-aware models that simulate tissue behavior, decode complex pathologies, and accelerate therapeutic innovation at unprecedented scale.

## INTRODUCTION

Human tissues normally maintain structural integrity and function across diverse physiological states, but how these architectures break down in disease, aging, and stress remains a fundamental, largely unresolved question in biology and medicine. The advent of spatial omics technologies has transformed our ability to map gene and protein expression within intact tissues, offering subcellular-resolution insights into cellular interactions and tissue architecture ^1,2^. Yet a critical gap remains: there is no comprehensive spatial reference of healthy human tissues across multiple organ systems. While existing atlases like the Human Cell Atlas ^3^, Human Protein Atlas, and Human Tumor Atlas Network ^4^, have advanced our understanding of single-cell biology, they are largely derived from dissociated cells or disease-specific samples, limiting their utility as healthy spatial baselines ^5^.

To address this critical need ^6^, we established the Spatial Atlas of Human Anatomy (SAHA)— the first multimodal, subcellular-resolution reference atlas of healthy adult human tissues spanning gastrointestinal and immune systems. SAHA integrates spatial transcriptomic, proteomic, and histological data from over 100 diverse donors, capturing cellular organization at ∼50 nm resolution. Unlike prior efforts, SAHA implements a rigorously standardized pipeline for tissue procurement, processing, quality control, and cross-platform integration. Our initiative enables consistent and reproducible analysis across organs, individuals, and spatial profiling technologies, while preserving the architectural context often lost in dissociated single-cell studies.

SAHA not only provides a high-resolution baseline of normal tissue organization but also facilitates comparative analyses with diseased tissues, enabling identification of spatially defined pathologies such as immune-epithelial remodeling, crypt dedifferentiation, and rare invasive niches. Through an open-access, FAIR-compliant ^7^ data portal (link to be updated upon acceptance), SAHA aims to serve as a foundational benchmark for translational research, computational modeling, and spatially informed diagnostics ^8^.

Here, we report the first phase of SAHA encompassing 16 gastrointestinal and immune tissue types and demonstrate its utility for spatially resolved analysis of diseases, such as colorectal cancer, pancreatic cancer, and inflammatory bowel disease. SAHA represents the largest subcellular resolution, multimodal transcriptomic and proteomic dataset of healthy human tissues to date, enabling the characterization of both conserved and organ-specific spatial niches. Through comparative analyses with diseased cohorts, we illustrate SAHA’s translational potential as a reference framework for identifying pathological alterations. By bridging spatial molecular profiling with clinical precision medicine, SAHA establishes a new paradigm for functional tissue atlas and a blueprint for the next generation of spatial omics research.

## RESULTS

### 1. Overview of the first gastrointestinal and immune spatial organ atlas

The first phase of the Spatial Atlas of Human Anatomy (SAHA) systematically mapped major digestive and immune organs at subcellular resolution, capturing over 15 million cells from over 100 healthy adult donors (ages 27-81 years, **Supplementary Table 1**). Using multiple high-plex spatial platforms including CosMx™ Spatial Molecular Imager, Xenium, RNAscope™, and GeoMx® Digital Spatial Profiler, we captured high-resolution profiles of stomach, liver, pancreas, small intestine (ileum), large intestine (descending colon and appendix), bone marrow, and lymph nodes. To provide a molecular bridge between spatially resolved tissue architecture and a more readily available clinical sample type, we also included peripheral blood mononuclear cells (PBMCs), which represent a non-solid tissue component (**Fig. 1, Supplementary Table 2**). We profiled 15,915,616 cells, including 9.5 million tissue-resident cells and 6 million circulating blood cells, at a spatial resolution of 50 nanometers. For the normal reference tissues, the CosMx RNA (1000-plex) dataset included 94 tissue samples, profiling 2.9 million cells and capturing over 524 million individual transcript measurements. This 1000-gene spatial atlas represents a substantial advancement in transcriptomic resolution— enabling detailed cell type phenotyping and cell–cell interaction analysis while preserving subcellular spatial fidelity. The corresponding CosMx protein dataset (67-plex) profiled 3.5 million cells and yielded over 17.8 billion quantified protein features. In contrast, disease and PBMC samples were analyzed using expanded RNA panels (6000-plex or 19,000-plex), providing deeper molecular coverage for characterizing rare populations and disease-specific states.

Standardized tissue collection and rigorous quality control were implemented to minimize batch effects and cross-platform variability, ensuring robust data integration and downstream analysis ^9^ (See **Methods**, **Extended Data Fig. 1, Supplementary Table 2**). This enabled joint analysis of transcriptomic profiles across organs, revealing both conserved and organ-specific cellular identities. Uniform Manifold Approximation and Projection (UMAP) of combined CosMx RNA data demonstrated distinct clustering of major epithelial lineages—such as hepatocytes (liver), acinar and ductal cells (pancreas), and gastric parietal and chief cells (stomach)—alongside broad immune populations including T cells, B cells, macrophages, and dendritic cells (**Fig. 2a, Extended Data Fig. 2a**). In total, over 50 distinct cell types were annotated using canonical marker genes, encompassing broad epithelial, immune, stromal, and neuronal lineages, highlighting both conserved and organ-specialized programs of cellular diversity and relative abundance (**Fig. 2b-d**).

Subcellular resolution enabled precise localization of transcripts and proteins within individual cellular compartments, allowing us to map the anatomical distribution of billions of biomolecules (RNA transcripts and proteins) across tissue sections. While higher resolution can increase technical variability and noise, our quality control strategy (i.e., histopathological review, quantification of control probes, RNAscope-based RNA integrity assessment, and platform cross-validation) ensured robust reproducibility across tissues and donors (**Extended Data Fig. 3**). For example, RNAscope control probes (i.e., *PPIB*) benchmarked RNA quality, and Spearman correlation analyses demonstrated a strong monotonic relationship (ρ = 0.86, p-value = 0.0007) between RNAscope and CosMx transcript measurements, with no detected systematic bias, while CosMx maintained greater transcriptome coverage due to its higher plex capacity (**Extended Data Fig. 3a-c**). Together, these analyses establish the technical reproducibility and biological validity of SAHA datasets and cell annotations, enabling robust cross-organ and cross-platform comparisons and providing a reliable reference for future spatial studies. Quality metrics across sample cohorts were highly consistent (**Extended Data Fig. 3d**), underscoring the robustness of the integrated data resource. This inaugural phase of SAHA establishes the largest high-quality, multimodal spatial reference of the human digestive and immune systems at subcellular resolution—providing a foundational resource for investigating tissue architecture, cellular diversity, and microenvironmental organization in health and disease.

### 2. Cellular neighborhoods and interactions across intestinal and immune structures

Leveraging an integrated embedding of over 15 million spatially profiled cells, SAHA enables both cross-organ comparisons and high-resolution analysis of spatial architecture within and across organ systems, tissue types, and cell populations (**Fig. 2**). This flexible framework supports multiscale investigation of tissue organization and microenvironmental interactions. In the gastrointestinal system, high-plex spatial profiling revealed nuanced, layer-specific architectures and discrete cellular niches spanning the epithelium and underlying submucosa (**Fig. 3a**). In the colon, for example, epithelial crypts were distinctly segmented, with *PIGR* and *MHC-I* expression localized to mucosal and submucosal regions, and smooth muscle markers (e.g., *ACTA2*) confined to the muscularis layer. Cell-level adjacency mapping further uncovered localized immune infiltration at crypt boundaries, suggesting active epithelial-immune crosstalk that is not readily detectable in dissociated single-cell datasets.

SAHA also provides a topographical link between structure and single-cell activity, wherein analysis of regionally enriched gene programs demonstrated spatial specialization along tissue depth axis (**Fig. 3b**). For example, outer crypt regions were enriched for keratinization-associated signatures, whereas deeper crypt zones expressed genes involved in antigen-processing and presentation (**Fig. 3b**). Spatial mapping of cell cycle regulators (e.g., *CDKN1A*, *CCND1*) and stress-response genes (e.g., *TFEB*) showed selective enrichment at immune-rich epithelial interfaces, suggesting localized zones of proliferative and immunomodulatory adaptation (**Fig. 3c**). Unbiased clustering of cell-cell adjacency networks further delineated distinct cellular neighborhoods across crypt and submucosal compartments (**Fig. 3d-f**). Discrete microenvironments dominated by epithelial cell to T cells, B cells, or myofibroblast interactions emerged along the crypt-luminal axis, emphasizing substantial spatial heterogeneity even within morphologically similar layers. These findings further highlight the unique ability of subcellular-resolution spatial profiling to uncover microanatomical, structural features that are inaccessible through dissociated single-cell approaches.

Beyond crypt-associated immune niches, gastrointestinal organs also revealed highly organized lymphoid structures. In the appendix (**Fig. 3g, Extended Data Fig. 4**), we identified large lymphoid aggregates composed of interspersed T and B cells surrounded by mesenchymal and myeloid populations. Force-directed graph visualizations of cell-cell adjacency networks demonstrated dense immune-immune interactions within these aggregates, consistent with their role as specialized hubs for adaptive immune activation (**Fig. 3h**). UMAP embeddings annotated by both broad and fine-grained cell identities highlighted tight spatial co-localization of functionally distinct immune subsets, including T follicular helper cells and germinal follicle center B cells (**Fig. 3i**). Ligand-receptor interaction analyses further revealed frequent paracrine signaling among these populations, suggesting coordinated immunological activity within these spatially confined niches (**Fig. 3j**).

These lymphoid aggregates occupy anatomically defined regions within gastrointestinal tissues and serve as sites of antigen presentation, lymphocyte activation, and localized immune coordination. To assess the generalizability of such spatially organized immune niches, we compared them with canonical germinal centers (GCs) in lymph nodes (LNs). Unsupervised clustering and spatial mapping of LN datasets revealed distinct compartmentalization between GC-resident B cells and surrounding immune and stromal populations (**Fig. 4a, Extended Data Fig. 5a-b**). Differential expression analysis further highlighted this organization, with B cell and antigen presentation genes (e.g., *CD74*, *CD79A, HLA-DRA*) enriched in GC cores (q-values = 5.47x10^-34^, 1.89x10^-10^, 1.20x10^-20^), while stromal and chemotactic transcripts (*CCL21, CXCL12, VIM*) localized to adjacent zones (q-values = 3.18x10^-99^, 7.21x10^-24^, 4.84x10^-20^) (**Extended Data Fig. 5c-f**).

To compare gut-associated lymphoid structures with canonical germinal centers, we analyzed anatomically paired epithelial (APE) lymphoid aggregates in the gastrointestinal tract. Similar to lymph node GCs, intra-GC B cells in APE structures expressed *IGHG1* and *IGKC*, while surrounding stromal regions were enriched for *APOE* and extracellular matrix genes (**Fig. 4c, Extended Data Fig. 5g**). However, unlike the sharply zonated architecture of lymph nodes, APE lymphoid structures were embedded within a more intermixed environment of epithelial and mesenchymal cells (**Fig. 4d-e**), suggesting greater potential for varied immune-epithelial crosstalk. Moreover, the SAHA data showed both generalized and tissue-specific transcriptomic gradients across GC and peri-GC regions in LN versus APE, highlighting organ-specific spatial organization (**Extended Data Fig. 5h**).

Expanding our analysis across all SAHA gastrointestinal tissues—including appendix (APE), ileum (ILE), colon (COL), and stomach (STO)—we profiled 43 patient samples comprising 948 fields of view (FOVs) and over 1.15 million spatially resolved cells (**Fig. 4g**). Integrative clustering of transcriptomic profiles combined with spatial adjacency information identified three major niche types: (1) organ-shared epithelial-immune interaction zones, (2) organ-specific immune microenvironments within lymphoid structures, and (3) rare cell populations with distinct spatial distributions (**Fig. 4h-k, Extended Data Fig. 6**). Notably, shared epithelial-immune neighborhoods, characterized by dense T- and B-cell infiltration adjacent to crypt structures, were recurrently observed in both colon and ileum samples (**Fig. 4h-i, Extended Data Fig. 6a- d**). In contrast, the stomach exhibited parietal-chief cell clusters unique to gastric glands, and the appendix contained specialized adaptive immune hubs, reflecting tissue-specific adaptations of immune-epithelial architecture (**Fig. 4j-k**).

To investigate the functional roles of these spatial niches, we leveraged the unique ability of spatial data to profile the ligand-receptor interaction networks between shared and organ-specific microenvironments. Conserved immune-epithelial niches were dominated by paracrine signaling between compartments, whereas tissue-specific structures exhibited distinct adaptive immune signaling profiles (**Extended Data Fig. 6e**). In addition, complementary spatial variance analysis revealed that genes involved in immune activation (e.g., *CD74*, *IGHM*), epithelial barrier integrity (e.g., *PIGR*), and chemokine signaling (e.g., *CCL21*) were highly localized within discrete spatial clusters (**Fig. 4l**). These findings suggest that spatial neighborhoods are defined not only by their cellular composition but also by localized transcriptional programs and interactions. Collectively, these analyses demonstrate that spatial information is essential for uncovering hierarchical tissue organization, functionally specialized microenvironments, and organ-specific immune architecture in the human gastrointestinal tract.

### 3. Validation and integration of multimodal spatial data

The detailed spatial mapping of gastrointestinal tissues highlights the power of subcellular-resolution atlases to uncover structured microenvironments and immune-epithelial interactions. To ensure the robustness and biological fidelity of these insights across platforms and tissues, we systematically validated SAHA datasets through multimodal integration of spatial transcriptomic and proteomic data. More importantly, we performed cross-platform validation using several orthogonal spatial assays, including CosMx RNA and protein profiling, 10x Genomics Xenium, RNAscope, GeoMx DSP, and histopathology-based annotations (**Supplementary Table 2**).

Integration of CosMx RNA and protein data enabled single-cell-level validation of biological consistency and facilitated deeper characterization of cell states and subtypes. Co-embedding of 1000-plex CosMx RNA and 67-plex CosMx protein profiles preserved tissue structure and cellular identities, confirming the robustness of multimodal spatial profiling (**Fig. 5a, Extended Data Fig. 7a-b**). Canonical markers such as CD45 (PTPRC), E-cadherin (CDH1), CD3D, and CD20 (MS4A1) exhibited concordant expression across RNA and protein modalities (**Fig. 5b-d**). Protein-based annotations aligned closely (Adjusted Rand Index, ARI = 0.2719) with RNA-integrated labels using Maxfuse ^10^, supporting consistent cell type classification across modalities and batches (**Fig. 5c, Extended Data Fig. 7c-d**). Protein data also provided clearer delineation of architectural features, such as the epithelial-stromal interface in the stomach and lymphoid aggregates in the appendix (**Fig. 5d**), reflecting the stability, localization, and abundance of structural proteins relative to transcriptomic signals ^11^. However, the transcriptomic profiles offered broader molecular coverage (up to 19k probes vs. <100 protein targets), enabling finer discrimination of epithelial subtypes and detection of rare populations such as stem and transit-amplifying cells.

Beyond validating cell identity and tissue structure, protein datasets offer orthogonal confirmation of receptor-ligand interactions inferred from RNA data (**Fig. 5e**) and provide direct evidence of translational activity. While such multimodal comparisons reinforce biological interpretation, it is important to note that RNA-protein correlations are not always linear, nor should it always be. Although this discordance has been examined for decades, systematic large-scale comparisons with subcellular resolution have remained limited ^12,13^. Leveraging SAHA’s multimodal and high resolution (50 nm), we directly compared matched RNA and protein measurements across multiple organs (**Fig. 5f**). Housekeeping and structural genes such as VIM showed strong concordance (adjusted R-squared of more than 0.5), whereas key immune and signaling molecules—including RELA and IGHD—displayed substantial discrepancies between RNA and protein levels (adjusted R-squared of less than 0.1) (**Fig. 5f, Extended Data Fig. 7e-f**). These differences were further highlighted in cross-organ analyses: VIM exhibited consistent RNA-protein correlation across tissues (R^2^ = 0.5781 with p-value = 0.0026 when compared across all SAHA cells), whereas IL1B showed widespread RNA expression with minimal protein detection. Conversely, CD276 demonstrated robust protein abundance despite low transcript levels, consistent with post-transcriptional regulation ^14^. These comparisons underscore the necessity of multimodal profiling to accurately capture both transcriptional potential and protein-level function.

The integration of histopathology-based annotations with spatial molecular profiles reinforced biological interpretability and enabled precise mapping of anatomical features to molecular states. Histological landmarks such as crypt architectures, lymphoid aggregates, and smooth muscle layers corresponded closely (qualitatively evaluated by cell types and niche clusters) labeled to spatial clustering patterns observed in molecular datasets (**Fig. 6**). This alignment between morphology and molecular identity validates the technical robustness and biological fidelity of the SAHA datasets, supporting consistent cross-tissue, cross-platform, and cross-modality analyses at subcellular resolution.

To further harness histological information, we applied CONCH, a multi-modal vision-text foundation model ^15^ that decodes tissue architecture from hematoxylin and eosin (H&E) stained whole-slide images (**Fig. 6a-b**). The model extracts quantitative morphological features from image tiles and learns spatial embeddings, which preserve anatomical continuity and discriminate between distinct structural zones (See **Methods**, **Fig. 6c-d**). Unsupervised (leiden) clustering of these image-derived features reconstructed tissue organization, including epithelial boundaries, stromal interfaces, and muscular layers (**Fig. 6e**). Visualization of representative clusters using UMAP revealed discrete and biologically meaningful groupings that corresponded to histologically distinct morphotypes, such as crypt bases, glandular epithelium, and lymphoid-rich regions most of which aligned with unbiased, RNA-based spatial clustering results (**Fig. 4g- k, 6f**).

This approach generalizes across gastrointestinal tissues, as image-derived embeddings from the colon, ileum, and appendix consistently showed conserved morphological classes, including crypt epithelium, extracellular matrix, and immune niches (**Fig. 6g**). Morphological clusters overlapped with specific histopathological terms scored by the multi-modal vision-text model (**Fig. 6h**) and displayed enrichment for specific tissue compartments such as epithelial, connective, or immune components (**Fig. 6i**). Spatial adjacency patterns among clusters (**Fig. 6j**) and their correlations with RNA-derived cell type proportions (**Fig. 6k**) further confirmed that image-derived features captured biologically relevant gradients in tissue organization. Finally, direct comparison of H&E image and CosMx RNA projection revealed highly consistent alignment shown by positive CCA correlation, between image-based morphology and transcriptomically defined cell types (**Fig. 6l**), illustrating the power of histology-informed molecular modeling for spatial atlas construction.

### 4. SAHA’s utility and impact in biomedical research

A key motivation for constructing SAHA was to enable comparative spatial analysis of diseased tissues, particularly in contexts where matched healthy controls are unavailable. To demonstrate its translational utility, we applied SAHA to two clinically relevant diseased conditions: colorectal cancer (CRC) and inflammatory bowel disease (IBD).

Using CRC samples, we analyzed transcriptional and structural profiles in tumor adjacent crypt regions and compared healthy tissue (SAHA COL, *n*L=L3) to histologically normal crypts from CRC resections (*n*L=L1) (**Fig. 7a**). Crypt regions were stratified and annotated as ’top’ and ’bottom’ crypts, based on gene expressions and their locations (**Fig. 7b, Extended Data Fig. 8a-b**). Bottom crypts were identified by enrichment of genes associated with intestinal stem cells (*OLFM4*) and differentiated secretory cells (*LEFTY1, STMN1*), whereas top regions expressed mature epithelial markers (e.g., *KRT20, PLAC8, CEACAM1*). Notably, crypts adjacent to CRC exhibited disruption in spatial expression patterns; markers normally restricted to bottom regions (i.e., *H2AZ1, SPINK1*) were aberrantly expressed in the upper crypts, and vice versa for *CEACAM6* and *CEACAM1* (**Fig. 7c, Supplementary Table 3**). These patterns potentially indicate crypt dedifferentiation and spatial disorganization often associated with tumor formation^16,17^.

Tumor-adjacent crypts showed widespread changes in gene expression, with significantly more genes up- or down-regulated compared to healthy crypts (**Fig. 7c**). Several upregulated genes, including *REG1A*, have been previously associated with poor prognosis and crypt-inflammation in CRC, while upregulation of *IL6* and *IL22* receptors are known to promote inflammation and tumor development in CRC ^18,19^. CRC-associated crypts had a substantially higher number of differentially expressed genes (DEGs) compared to COL crypts, indicating broader transcriptional reprogramming (i.e., 96 vs 61 DEGs in bottom crypt and 116 vs. 72 DEGs in top crypt; **Supplementary Table 3**). In terms of the tumor microenvironment, FOVs profiled from CRC samples showed a significant increase in the number of mast cells (**Extended Fig. 9c**), a known driver of angiogenesis and metastasis ^20^.

To further investigate spatial expression and organization of RNA transcripts, we performed spatial autocorrelation analysis using Moran’s I and clustered FOVs based on their top 200 spatially variable genes (SVGs) (**Fig. 7d**). This unsupervised approach effectively distinguished crypts from CRC and COL samples, indicating their divergent spatial states. From matched top and bottom crypt compartments across conditions, we visualized all SVGs and identified both conserved spatial markers (e.g., *KRT20, CEACAM1, CEACAM6*) and spatially enriched disease-associated genes (e.g., *REG1A, IL22RA1*) - both of which were also noted in our differential expression analysis (**Fig. 7e-f, Extended Data Fig. 8e**).

Beyond structural comparisons, SAHA enabled exploration of intrapatient tumor heterogeneity (**Fig. 7g-j**), including rare events such as perineural invasion (PNI) (**Fig. 7k-m**). Within a single CRC sample, we identified two spatially and transcriptionally distinct tumor subtypes (labelled as Type 1 and Type 2), based on dominant cell compositions and immunogenicity (**Fig. 7g**).

Type 1 tumor regions exhibited a higher lymphocyte to tumor ratio with increased proportions of cytotoxic and naive CD8+ T cells, whereas Type 2 regions were enriched for macrophage subsets (**Fig. 7g-j, Extended Data Fig. 8f-g**).

We captured a rare perineural invasion event among the analyzed FOVs, characterized by dysplastic stroma surrounding a nerve and tumor cells encircling the perineurium (**Fig. 7k**). Although perineural invasion is recognized as a hallmark of aggressive CRC, it remains difficult to resolve with conventional histopathology and immunofluorescence, which lack comprehensive transcriptomic resolution ^21–23^ . While recent single-cell RNA-seq studies have attempted to profile these events, they often fail to capture neural cells altogether—or, if captured, cannot confidently localize them or distinguish whether they are part of a PNI structure or simply present in unrelated regions of the tissue ^24–26^. By leveraging the spatial context of SAHA, we subclustered this PNI region and uncovered a rare subset of fibroblasts (less than 100 cells in the PNI region) in close proximity to glial cells, characterized by a hybrid transcriptional signature expressing both glial-associated genes (*CLU, NGFR, SLC2A1, APOD*) and cancer-associated fibroblast markers (<2% of fibroblasts identified in the sample) such as *CXCL14* and *MMP2* ^27,28^ (**Fig. 7k-l, Extended Data Fig. 8h**). Cell-cell communication analysis further revealed unique signaling pathways within the PNI niche, including the ANGPTL pathway originating from fibroblasts, previously associated with increased metastatic risk in CRC ^29–31^, and the LIFR pathway, exclusively signaling from tumor cells to glial cells, known to facilitate tumor-fibroblast crosstalk and activate pro-invasive programs ^32^. These findings offer new insights into the interactive dynamics within this rare microenvironment and its ability to specify high resolution details of rare tumor types.

We also compared SAHA ILE with ileal samples from patients with inflammatory bowel disease (IBD), including individuals stratified by clinical response to TNFα inhibitor therapy (**Fig. 8a**).

Integration of CosMx RNA profiles across healthy and IBD ileum samples revealed clear separation by disease state, while preserving major cell type identities (**Fig. 8b, Extended Data Fig. 9a-b**). Correlation analysis of average gene expression profiles demonstrated strong consistency (e.g., spearman correlation R^2^ of 0.74 for epithelial cells from 1K and 6K datasets) across datasets generated with different plex sizes, validating the reproducibility of cell-type transcriptional programs (**Extended Data Fig. 9a**). Spatial mapping of cell types demonstrated increased immune cell infiltration into epithelial compartments in IBD samples relative to healthy controls (**Fig. 8c, Extended Data Fig. 9c-d**).

Quantification of epithelial-proximal immune infiltration within 300 μm confirmed a significant increase in immune cell density in IBD tissues (p < 0.001; **Fig. 8d**). Ligand–receptor interaction analyses further revealed enhanced immune-epithelial signaling in IBD, particularly among non-responders to TNFα therapy, who exhibited elevated spatial interaction scores relative to responders (**Fig. 8e**). Mesenchymal-stromal populations in IBD tissues showed increased expression of pro-inflammatory and extracellular matrix remodeling markers, including elevated *TNFRSF1A* (TNF receptor 1) expression among non-responders compared to responders (p < 0.05; **Fig. 8f**). Spatial mapping of TNF-TNFRSF1A ligand–receptor interactions localized active signaling to immune-mesenchymal interfaces, particularly in non-responder tissues (**Fig. 8g**).

Moreover, these molecular level changes were consistent with the histological analyses. As performed for the healthy GI tract (**Fig. 6**), we applied a multi-modal text-vision model to quantify and describe the morphological patterns of H&E-stained ileum sections of healthy and IBD tissues (**Fig. 8h**). IBD samples were distinguishable from healthy, with histopathological terms related to metaplasia, immune cell infiltration, crypt branching, and apoptosis more frequent in IBD samples compared to controls (**Fig. 8i**). Furthermore, we also found differentially enriched morphological changes in response to TNFα treatment (**Fig. 8i**), which were aligned with molecular findings, reinforcing the multi-layered disruption of epithelial and stromal architecture in IBD ^33^. These analyses demonstrate how SAHA enables spatially resolved comparisons of healthy and diseased tissues at both molecular and histological levels, providing critical insights into disease-associated microenvironmental changes and therapeutic response heterogeneity.

The integration of spatial transcriptomics and proteomics extends the analytical frameworks traditionally applied to single-cell sequencing by adding critical layers of structural and contextual information (**Fig. 9a**). Spatial multi-omics enables not only transcriptomic and proteomic profiling at subcellular resolution, but also the characterization of extracellular matrix (ECM) features, direct cell-cell interactions, and subcellular biomolecule localization, which are dimensions inaccessible to dissociated single-cell methods. The addition of spatial context enables direct measurement of cellular phenotypes, resolving the challenge of drawing conclusions from disconnected data. This multidimensional mapping redefines efforts in building biological atlases and opens new avenues for understanding tissue organization and function.

Recent technological advances now enable high-throughput, same-cell spatial multi-omics at subcellular resolution—capturing whole-transcriptome RNA (∼19,000 genes), high-plex protein expression (∼67 markers), and histological context within a single assay (**Fig. 9a**). These spatial platforms also provide cell morphology, boundary, and segmentation data that are inaccessible via traditional dissociated single-cell methods. To scale this approach across large tissue volumes and billions of cells, we implemented computational workflows capable of integrating transcriptomic, proteomic, and H&E morphological data at population scale. Compared to conventional scRNA-seq, spatial multi-omics approaches using Xenium and CosMx offer orders-of-magnitude lower cost per cell and support efficient scaling to billions of cells with fewer experimental runs—facilitating population-scale spatial atlasing (**Fig. 9b**).

To maximize accessibility and impact, SAHA adheres to the FAIR (Findable, Accessible, Interoperable, and Reusable) data principles. The SAHA data portal provides an open-access platform for exploring spatial multi-omics datasets, equipped with interactive visualization tools, high-resolution tissue navigation, and download capabilities for downstream analysis (**Fig. 9c**). Beyond serving as a reference, SAHA establishes standardized protocols and integration frameworks that can be adopted across institutions to enable scalable, reproducible spatial profiling.

## DISCUSSION

The Spatial Atlas of Human Anatomy (SAHA) establishes the first multi-organ, multimodal, and multi-platform reference map of healthy human gastrointestinal and immune systems at subcellular resolution, providing a foundational resource for investigating tissue architecture, cellular diversity, and microenvironmental organization for healthy and diseased states. By integrating histological, proteomic, and transcriptomic data across tissues and spatial technologies, SAHA addresses major limitations of previous atlasing efforts, which have predominantly relied on dissociated single-cell data and lacked spatial context. A key strength of SAHA lies in its rigorously standardized experimental and computational pipelines, which ensure reproducible, high-quality data generation and facilitate integrative analyses across organs and modalities. To support cross-platform consistency, we implemented systematic validation across CosMx, Xenium, GeoMx, and RNAscope platforms, alongside robust quality control metrics. While SAHA initially leveraged 1000-plex RNA profiling, we also introduce, to our knowledge, the first whole-transcriptome and paired protein multi-omics datasets at subcellular resolution—expanding beyond mid-plex spatial transcriptomics to unlock deeper biological insights and establish a future-ready reference for spatial biology.

A central innovation of SAHA lies in its multimodal design and backwards (H&E)-and-forwards (full transcriptome) compatible framework. The integration of RNA, protein, and histopathological (H&E) data facilitates both cross-validation of molecular signals and nuanced dissection of tissue structure and function. This multimodal synergy enabled identification of conserved spatial niches, lineage-specific expression gradients, and regionally restricted cell states, including rare and adaptive immune populations such as follicular B cells and intraepithelial lymphocytes. We also uncovered fine-grained epithelial and stromal subtypes, such as tuft and enterochromaffin-like cells, and delineated spatially encoded state transitions within common lineages—patterns that would be obscured in single-modality or dissociated datasets. Some segmentation ambiguity is inevitable in a subset of cells, in part due to the physical limitations of tissue sectioning through three-dimensional structures. However, we observe that such instances actually encode meaningful biological information, as spatial proximity—retained in intact tissue—offers a critical dimension absent in single-cell suspensions. Dissociation-based workflows may also lose spatially tethered subpopulations during washing and library preparation steps. While future pixel-level or nucleus-aware segmentation methods ^34,35^ may refine these boundaries, our current analysis emphasizes broader tissue-level organization, highlighting reproducible spatial relationships that extend beyond cell-intrinsic expression.

Across gastrointestinal tissues—including appendix, ileum, colon, and stomach—SAHA captures a conserved layered organization spanning mucosa, submucosa, muscularis externa, and serosa. At the same time, SAHA reveals organ-specific features, such as crypt-villus gradients, lymphoid aggregates (e.g., Peyer’s patches), and vascular-rich stromal zones. These features were consistent across individuals and histologically validated, reinforcing the atlas’s technical fidelity. Notably, cross-organ analysis within gastrointestinal tissues uncovered conserved features of mucosal organization while highlighting unique adaptations in the appendix, ileum, colon, and stomach. A comparative profiling of lymphoid aggregates across GI organs and matched germinal centers in lymph nodes revealed both preserved core immune architectures and distinct spatial intermingling with epithelial and stromal compartments, emphasizing the utility of SAHA in decoding tissue-specific immunological microenvironments.

Importantly, SAHA serves not only as a reference but also as an actionable spatial comparator for disease studies—especially in clinical contexts where matched healthy tissue is unavailable. We demonstrate this translational potential by benchmarking SAHA against datasets from colorectal cancer (CRC) and inflammatory bowel disease (IBD). This revealed distinct alterations in crypt top–bottom zonation, immune composition, and spatial gene expression programs. For instance, spatial transcriptomics exposed crypt dedifferentiation in CRC and aberrant cytokine signaling, while perineural invasion—an aggressive metastatic route—was resolved at high granularity, highlighting glia-associated fibroblast states and tumor-nerve signaling circuits. These examples illustrate how SAHA enables spatial biomarker discovery and mechanistic insight into rare or complex pathological events, areas that have been briefly explored in previous studies.

While SAHA represents a significant advance, several limitations remain. First, despite including over 100 donors, our cohort size is modest relative to the full scope of human biological diversity. Further expanding SAHA across age groups, ancestries, and environmental exposures is essential for broader generalizability. Second, although cross-platform integration across CosMx, Xenium, GeoMx, and RNAscope demonstrates strong concordance, inherent technical differences—such as probe capture efficiencies and antibody variability—necessitate ongoing methodological harmonization. Third, this initial phase focuses on gastrointestinal and lymphoid organs; a complete human reference will require comprehensive coverage across additional anatomical systems.

While the current datasets represent the initial phases of spatial atlas construction, the platform is designed for scalability and community-driven expansion. Active efforts are underway to expand its organ coverage (second phase to genitourinary systems and additional immune organs), apply whole-transcriptome and high-plex proteomics protocols, increase demographic representation, and integrate advanced computational frameworks, including deep learning for tissue segmentation and spatial graph modeling. We also aim to leverage Common Coordinate Framework (CCF)-based registration and analysis pipelines to enable 3D spatial integration across tissue sections and donors. This will support more anatomically precise mapping, facilitate multi-plane reconstruction of tissue architectures, and allow in-depth cross-sample comparisons at scale.

By integrating spatial molecular data with histological and clinical metadata, SAHA provides the infrastructure for building digital twins of human tissues—enabling in silico simulations of disease progression, therapeutic response, cell-cell interactions, and possibly even tissue regeneration. These capabilities position SAHA as a translational bridge between spatial biology and precision medicine, advancing both fundamental research and clinical diagnostics. For example, we can extend SAHA toward perturbation-aware modeling, using spatial benchmarks to assess disease-associated changes and therapeutic responses—analogous to recent large-scale single-cell initiatives such as Tahoe-100M ^36^. Together, these directions will transform SAHA into a foundational tool for spatially resolved systems biology and experimental medicine.

The SAHA data portal, built on FAIR (Findable, Accessible, Interoperable, Reusable) principles, will enable broad community access and invite collaborative expansion. We envision that SAHA will not only support basic biological discovery and accelerate clinical translation by providing a standardized framework for the development of spatially informed diagnostics, prognostics, and therapies. As spatial omics technologies continue to evolve, SAHA serves as a dynamic foundation to integrate emerging data types, drive systems-level insights into tissue function and disease, and chart new frontiers in precision medicine. We invite researchers worldwide to utilize SAHA datasets, develop new analytical tools, and contribute additional spatial multi-omics data to collectively accelerate biological discovery and the construction of a comprehensive human spatial atlas.

The integration of SAHA with artificial intelligence frameworks will catalyze a new generation of spatially aware models capable of learning complex tissue organization, molecular gradients, and context-specific cell-cell interactions ^37,38^. Unlike dissociated single-cell datasets, SAHA’s subcellular-resolution, multimodal profiles retain anatomical fidelity, enabling training of deep learning models that incorporate both spatial topologies, molecular features, and next generation omics. This includes the development of graph neural networks for cell-cell communication inference, foundation models for cross-tissue pattern recognition, and generative architectures for simulating disease progression or therapeutic response within intact tissue contexts. Moreover, SAHA’s structured metadata, ontology-linked annotations, and standardized formats (e.g., AnnData ^39^, OME-TIFF ^40^) make it inherently machine-readable and interoperable across AI pipelines. By coupling biological fidelity with computational scalability, SAHA provides a foundational resource for training AI systems capable of performing high-resolution tissue segmentation, phenotype classification, and predictive modeling across spatial, molecular, and morphological domains.

## METHODS

### Patient material, ethics approval, and consent for publication

Human tissue samples were obtained from archived formalin-fixed, paraffin-embedded (FFPE) blocks curated by board-certified pathologists and collaborating investigators (**Supplementary Table 1**). Each tissue sample was selected from a diverse ancestry (if possible) from healthy adults (no cancer or organ wide immune issues, ideal age 20-40). If tissues were difficult to get, age and ancestry parameters can be compromised, but no cancer patients or patients with diseases that will cause an impact on not diseased tissues was considered. Representative fields of view (FOVs) were selected following comprehensive histopathological evaluation of H&E-stained slides. All samples were collected under pathology protocols with waivers of consent and authorization.

Tissue microarray (TMA) formalin-fixed, paraffin-embedded (FFPE) blocks were constructed following standard procedures. SAHA FFPE blocks were selected based on histopathological evaluation, and representative areas were identified on hematoxylin and eosin (H&E)-stained slides. Tissue cores (variable in diameter depending on the organ) were extracted using a manual or semi-automated tissue arrayer and transferred into a pre-designed recipient paraffin block. The TMA block was re-embedded and cooled to ensure structural integrity. SAHA FFPE blocks were stored in cardboard boxes at room temperature (RT) under ambient humidity until the day of sectioning.

### Tissue processing and histology

The FFPE blocks were sectioned at a thickness of 5 µm, and the sections were placed within Bruker’s 15mm x 20mm allowed scan area on Leica Bond Plus slides (Leica Biosystems, S21.2113.A). Fresh ultrapure water was used to fill the water bath and new microtome blades were used for each tissue type. The first few sections from the block face were discarded. The number of serial sections to be collected was determined in advance according to the planned assays. Slides were labeled with a serial number to keep track of the order, ensuring that multiple assays were run on adjacent sections and facilitate subsequent data integration. If needed, one section, usually from the middle of the series, was dedicated to H&E staining, followed by full-slide image acquisition using the Aperio Digital Pathology Slide Scanner system (Leica Biosystems), allowing pathologists to identify and mark the area of interest in consideration of the available number of CosMx fields of view (FOVs). Most sections were also H&E stained after data acquisition. **Supplementary Table 2** lists the serial section numbers for each assay and for the corresponding H&E for each analyzed organ. Following sectioning, the slides were dried upright at RT for 30 minutes (min), followed by a horizontal overnight (ON) drying step at 37°C in an oven to improve tissue adherence. The slides were then stored in a desiccator at RT until processing. Most slides dedicated to the RNA assay were used within two weeks, as recommended by Bruker. Exact sectioning and run dates are reported in Supplementary Table 2.

### CosMx processing

Processing of the slides followed the guidelines outlined in the Bruker manuals MAN-10184 for the RNA assay and MAN-10185 for the Protein assay, utilizing the equipment, materials, and reagents recommended in the manuals. Protocol details may have varied over time, reflecting updated manuals throughout the SAHA project. The specific manual version used for each assay is reported in **Supplementary Table 2**.

For the CosMx RNA assay, slides were baked at 60°C ON in vertical position in a baking oven. The following day, CitriSolv (VWR, 89426-268) was used for deparaffinization. Specifically, slides were brought to RT for 3 min, then subjected to two sequential 5 min immersions in CitriSolv, followed by two 5 min washes in 100% ethanol (Et-OH) (Sigma, E7023). The slides were dried vertically for 5 min at 60°C in the same baking oven. Antigen retrieval was conducted in a BioSB® TintoRetriever digital pressure cooker (BioSB, BSB 7015) at 100°C for 15 minutes. After the 5-minute drying step, slides were transferred into the preheated 1X Target Retrieval Solution (part of Bruker 121500006) as per the Bruker manual. The pressure cooker was then closed, and the temperature was allowed to return to 100°C. At this point, a 15 min timer was started, after which the slides were quickly rinsed up and down in DEPC water (Fisher Scientific, BP561-1) at RT for 15 seconds, washed for 3 min in 100% Et-OH, and then dried for 30 min horizontally on a clean surface at RT. 10 min before the end of the drying step, the incubation frame (part of Bruker 121500006) was applied to the slide and a 3 µg/mL Proteinase K (part of Bruker 121500006) working solution in 1XPBS (ThermoFisher Scientific, AM9625) was applied to the sections. The slides were then incubated for 30 min at 40°C in a RapidFISH slide hybridizer oven (Boekel Scientific, 240200). During the final 10 min of digestion, Bruker fiducial beads (part of Bruker 121500006) were resuspended through alternating 1-min vortexing and 2- min sonication steps (three vortexing steps and two sonication steps in total), then diluted in 2X SCC-T (part of Bruker 121500006). Fiducials were diluted at a final working concentration of 0.001% for Appendix, Ileum, Colon, and Pancreas samples, 0.00075% for Stomach and Lymph Nodes samples, and 0.0005% for inflammatory bowel disease ileum samples. Following digestion, slides were washed twice in DEPC water per the manual and then incubated with the prepared fiducials for 5 min, protected from light at RT. The slides were then washed in 1XPBS for 1 min, fixed in 10% Neutral Buffered Formalin (NBF) (EMS Diasum, 15740) for 1 min, washed twice in a 0.1M Tris-Glycine NBF buffer solution for 5 min, once in 1XPBS for 5 min, and incubated with a freshly prepared 100 mM NHS-Acetate (Fisher Scientific, 26777) solution in NHS-Acetate buffer (part of Bruker 121500006) for 15 min in the dark at RT. Finally, the slides were washed twice in 2X SSC (ThermoFisher Scientific, AM9763) for 5 min. At this point, slides were stored in 2X SSC, protected from light, until it was possible to proceed with the ON hybridization step. The required amount of probe for the run was transferred to 0.2 mL tubes, denatured for 2 min at 95°C in a standard thermocycler, and crash-cooled on ice for 1 min. A probe hybridization mix containing RNase inhibitors (Bruker, 121500004) and Buffer R (part of Bruker 121500006) was prepared according to the Bruker manual. Slides were laid horizontally on a clean surface and the hybridization mix was applied to the sections. A plastic coverslip (part of Bruker 121500006) was then placed over the frame. The slides were incubated ON for 17-18 hours (h) at 37°C in the RapidFISH slide hybridizer oven. The RNA probe panel used for the SAHA project was the CosMx™ Human Universal Cell Characterization Panel (Bruker, 121500002), except for inflammatory bowel disease ileum samples, which were run using the CosMx™ Human 6K Discovery Panel (Bruker, 121500041). On the following day, after removing the coverslip, slides were quickly rinsed in 2X SCC, washed twice for 25 min at 37°C in pre-warmed 50% 4X SSC/50% formamide (ThermoFisher, AM9342), and washed again in 2X SCC twice for 2 min. The slides were then incubated in the dark at RT for 15 min with a 1:40 dilution of DAPI (part of Bruker, 121500020) in blocking buffer (part of Bruker, 121500006). After a 5-min wash in 1XPBS, cell segmentation marker staining was performed using a cocktail of CD298/B2M, PanCK, CD45, and CD68 antibodies. The antibodies are conjugated to a readout domain that binds fluorophores on-instrument. Slides were incubated in the dark at RT for 1 h with the morphology staining marker mix, prepared using the CosMx™ Human Universal Cell Segmentation Kit, RNA (Bruker, 121500020) and the CosMx™ Human CD68 A La Carte Marker, RNA (Bruker, 121500022), at the dilutions specified in the Bruker manual. Slides were then washed three times in 1X PBS for 5 min and transferred to 2X SSC. For most samples, the protocol spanned three days, meaning that at this stage, the adhesive frame was removed, and slides were stored ON in 2X SSC at 4°C, protected from light. The only exceptions were stomach and lymph node samples, which were fully processed and loaded on the instrument in just two days. On the following day, the flow cells were assembled using the Flow Cell Assembly Tool provided by Bruker and Bruker glass coverslips with ports (CosMx™ Flow Cell Assembly Kit, Bruker, 122000061). They were then kept in 2X SSC, ensuring the solution filled the flow cells to prevent section dehydration, and stored in the dark until loading onto the CosMx instrument.

For the CosMx Protein assay, slides were baked for just 1 h at 65°C in a vertical position.

Similar to the RNA assay, after baking slides were equilibrated for 3 min at RT and then immersed in CitriSolv twice for 5 minutes. In the case of the protein workflow, this was followed by a series of washes: two 10 min washes in 100% Et-OH, two 5 min washes in 95% EtOH, one 5 min wash in 70% EtOH, and two 5 min washes in 1XPBS. The target retrieval step was performed as described above for the RNA assay, with the only difference that after the 15 min retrieval, the solution containing the slides was left to cool at RT for 25 min. The slides were then transferred to 1XPBS and washed three times for 5 min. At this point, the incubation frame (part of Bruker 121500008) was applied to the slides. Sections were incubated with blocking Buffer W (part of Bruker 121500008) for 1 h at RT, followed by ON incubation at 4°C with a single mix containing both the morphology marker antibodies for segmentation (CosMx Human Universal Cell Segmentation Kit, IO PanCK/CD45 Supplemental Segmentation Kit, and CD3 A La Carte Marker, Bruker 121500026, 121500027, and 121500028) and the antibodies from the CosMx Human Immuno-oncology 64-plex panel (Bruker, 121500010). The sections were covered with a plastic coverslip (part of Bruker 121500008) during ON incubation. The following day, after removing the coverslip, the slides were washed three times for 10 min in 1XTBS-T (ThermoFisher, J77500.K2) and once for 2 min in 1XPBS. During the 1XTBS-T washes, fiducials were prepared as for the RNA assay. For the protein assay, the final fiducial working concentration was 0.00005%, requiring a two-step serial dilution in 1XTBS-T, as recommended by the manual. After a 5 min incubation at RT with fiducials, slides were washed once with 1XPBS for 5 min, fixed with 4% PFA (Electron Microscopy Sciences, 15710) for 15 min at RT in the dark, washed three times in 1XPBS for 5 min each, and incubated for 10 min at RT with a 1:40 dilution of DAPI (Bruker 121500026) in 1XPBS. The slides were then washed twice more in 1XPBS for 5 min, incubated with freshly prepared 100 mM NHS-Acetate for 15 min protected from light at RT, and subjected to a final 5 min wash in 1XPBS. As for the RNA assay, for most of the SAHA protein assays, the protocol extended over three days. Slides were transferred to fresh 1XPBS tubes and stored at 4°C in the dark ON. On the following day, flow cells were assembled as described above, kept in 1XPBS in the dark, and loaded onto the CosMx instrument.

To set up a new CosMx runs on the Bruker CosMx Spatial Molecular Imager (SMI), the Bruker manual MAN-10161 was used as a guide. Different versions of the manual were followed throughout the SAHA project, and various SMI instrument software versions were installed on the machine over time. **Supplementary Table 2** lists the instrument manual and software versions for each analyzed sample. The SMI Control Center webpage interface allows for the setup of all run specifications, as well as the assignment of FOVs. The setup begins with configuring a new flow cell, the first step of which is defining the primary information for the run, including the Pre-Bleaching Profile and the Cell Segmentation Profile. As these profiles are tissue-type specific, they were selected based on Bruker’s recommendations in MAN-10161, but they ultimately ended up being Pre-Bleaching Profile C and Cell Segmentation Profile A for all our runs, both RNA and protein. The probe panel and cell segmentation kits used also needed to be specified at this stage. The sensor-based interactive SMI Control Center interface guided the loading of the instrument. The Imaging Tray (CosMx RNA Imaging Tray, Bruker 122000156, 122000157, or 122000158, CosMx Protein Imaging Tray, Bruker 122000162 or 122000163), flow cells, and buffers required for the run (CosMx RNA Instrument Buffer Kit, Bruker 100480 or 100481, CosMx Protein Instrument Buffer Kit, Bruker 100482) were loaded into their designated slots. For RNA assays, RNase inhibitors (Bruker, 121500004) were added to a specific well of the Imaging Tray. For all runs, two enzymes - Catalase and Pyranose Oxidase (part of Bruker 100480, 100481, and 100482) - needed to be resuspended, centrifuged, and added free of precipitate to Buffer 4 in advance. Once the assembled flow cell were loaded onto the instrument and all run parameters were approved, the system proceeded through a deck validation step, an instrument preparation step, and a preliminary morphology marker scan to allow the FOVs placement. RNA target readout on the CosMx SMI instrument was performed as described in He et al. ^41^. Reporter Wash, Imaging, and Strip Wash Buffers all supplied by Bruker Spatial Biology. At this point, the FOV Selection Workspace opened, and the FOV Placement Tool enabled for the placement, movement, or deletion of individual FOVs or grids of FOVs over the scanned image of the section. For large tissue areas, a serial H&E scan was used in advance by pathologists to annotate the image and define the regions of interest to be covered by the available FOVs. For TMAs, we aimed to allocate a predefined number of FOVs to individual tissue cores. Once the FOVs were reviewed and approved, the machine began cycling. RNA readout began by flowing 100 μl of Reporter Pool 1 into the flow cell and incubating for 15 min. Reporter Wash Buffer (1 mL) was flowed to wash unbound reporter probes, and Imaging Buffer was added to the flow cell for imaging. Eight Z-stack images (0.8 μm step size) for each FOV were acquired, and photocleavable linkers on the fluorophores of the reporter probes were released by UV illumination and washed with Strip Wash buffer. The fluidic and imaging procedure was repeated for all the reporter pools, and the different rounds of reporter hybridization (based on assay plex) imaging were repeated multiple times to increase target detection sensitivity. Cell morphology was imaged on-instrument prior to RNA or protein readout by adding fluorophore-bound reporters to the flow cell and capturing eight z-stack images in the channels 488 nm (CD298/B2M), 532 nm (PanCK), 594 nm (CD45), 647 nm (CD68), and 385 nm (DAPI). Runs lasted anywhere from two to ten days, depending on the panel type and plex, and the number of FOVs. CosMx data were automatically uploaded to Bruker’s cloud-based AtoMx Spatial Analysis Platform both during and after the run.

### Single-nuclei sequencing (snPATHO-seq)

Single nucleus RNA sequencing (snRNA-seq) was performed following the snPATHO-Seq protocol as previously described ^42^. Two scrolls of 30–50 µm thickness were obtained from each FFPE sample, adjacent to the sections used for CosMx, Xenium, and H&E staining. Scrolls were collected into 1.5-mL Eppendorf tubes, stored at 4°C in Ziplock bags containing desiccants to control humidity, and processed within one week. For nuclei isolation, scrolls were dewaxed by three sequential washes in 1 mL of xylene (Millipore Sigma, 214736) for 10 minutes each, with the first xylene wash performed at 55°C to enhance paraffin removal. Rehydration was achieved through serial immersions in decreasing concentrations of ethanol (100% twice, followed by 70%, 50%, and 30%; each for 1 minute) (Millipore Sigma, E7023), followed by a single wash with 1 mL of RPMI 1640 medium (Gibco, 11875093).

Tissues were then digested in RPMI 1640 supplemented with 1 mg/mL Liberase TH (Millipore Sigma, 5401151001) and 1 U/µL RNase inhibitor (Thermo Fisher Scientific, EO0382). The scrolls were fragmented using a pellet pestle, and the digestion volume was adjusted to 1 mL. Digestion proceeded at 800 rpm in a Thermomixer. Following digestion, 400 µL of EZ Lysis Buffer (Sigma-Aldrich, NUC-101) was added, mixed, and centrifuged at 850 × g for 5 minutes at 4°C. The nuclei-containing pellet was resuspended in EZ Lysis Buffer with 2% BSA (Miltenyi, 130-091-376) and RNase inhibitor, then homogenized by pestle and pipetting. The suspension was filtered through a 70-µm strainer (pluriSelect, 43-10070-50) and centrifuged. The nuclei pellet was washed three times with 1× PBS containing 1% BSA (Gibco, 70011044), followed by two washes with 0.5× PBS containing 0.02% BSA. Final resuspension was performed in 500– 1000 µL of 0.5× PBS with 0.02% BSA, depending on pellet size, and filtered through a 40-µm strainer (pluriSelect, 43-10040-50).

Nuclei were quantified using the Luna-FX7 instrument with Acridine Orange/Propidium Iodide staining (Logos). For immediate processing, nuclei suspensions were kept on wet ice. snRNA-seq libraries were prepared using the Chromium X Controller (10x Genomics) and the Chromium Fixed RNA Profiling Reagent Kits for multiplexed samples (Chromium Fixed RNA Kit, PN-1000475; Chromium Next GEM Chip Q, PN-1000418; Dual Index Kit TS Set A, PN- 1000251), following the manufacturer’s user guide (CG000527, Rev E). For cryopreservation, nuclei suspensions were supplemented with 10% glycerol and 0.1× Enhancer (10x Genomics, PN-2000482), rested on ice for 10 minutes, and stored at -80°C.

Hybridization was performed using 500,000–1.5 million nuclei per sample at 42°C for 20 hours. Barcoding reagents BC1–BC4 were used. Post-hybridization washes were conducted in a pooled setting according to the user guide with an additional wash step. Approximately 10,000 nuclei were targeted for capture per sample. An extra cycle was added during the pre-amplification PCR step, and indexing PCR was optimized to achieve final library concentrations between 50 and 200 nM. Library quality was assessed using a Fragment Analyzer 5200 (Agilent) and quantified using a Qubit Flex Fluorometer (Thermo Fisher Scientific). Libraries were sequenced on an Illumina NovaSeq 6000 using a 28 × 10 × 10 × 90 bp read configuration, targeting 20,000–30,000 read pairs per nucleus.

Transcript count matrices generated by Cell Ranger v8.0.0 (10x Genomics) were imported into Seurat v5.0.1 for downstream analysis ^43–47^. Doublet identification was performed using scDblFinder v1.14.0 with a preset doublet rate of 10%, reflecting the expected frequency in Single Cell Gene Expression Flex multiplexing experiments ^48^. Quality control based on UMI and gene count distributions identified a population of low-complexity droplets, which lacked distinct transcriptomic signatures and were excluded from further analysis. Nuclei flagged as doublets or containing fewer than 1,000 detected genes were removed, ensuring retention of major cell populations as validated by comparison with the original dataset. Normalization and highly variable feature selection were conducted using SCTransform with default parameters. Dimensionality reduction was performed using principal component analysis (PCA), with the first 30 principal components guiding Uniform Manifold Approximation and Projection (UMAP) embedding (RunUMAP) and Louvain clustering.

### Xenium Slide Preparation and In Situ Gene Expression Assay

Sections for Xenium assays were prepared according to the *Xenium In Situ for FFPE—Tissue Preparation Guide* (CG000578 Rev C, 10x Genomics) following the manufacturer’s instructions. Briefly, 5-µm-thick FFPE tissue sections were floated in an RNase-free water bath and mounted onto the 10.45 mm × 22.45 mm sample area of Xenium slides (PN-3000941, 10x Genomics).

Slides were air-dried at room temperature for 30 minutes, followed by a 3-hour incubation at 42°C on a Xenium Thermocycler Adapter plate positioned atop a 96-well thermal cycler (C1000 Touch, Bio-Rad) with the lid open. Slides were further dried ON at room temperature in the presence of desiccants.

Subsequent processing, including deparaffinization and decrosslinking, was performed following the *Xenium In Situ for FFPE—Deparaffinization and Decrosslinking Protocol* (CG000580 Rev C, 10x Genomics). Slides were incubated at 60°C for 2 hours, equilibrated at room temperature, then sequentially immersed in xylene (twice, 10 minutes each), 100% ethanol (twice, 3 minutes each), 96% ethanol (twice, 3 minutes each), 70% ethanol (once, 3 minutes), and water (once, 20 seconds). Following rehydration, slides were assembled into Xenium cassettes and hydrated with 1× PBS. Tissue sections were reverse crosslinked using a decrosslinking buffer containing tissue enhancer, urea, and perm enzyme B at 80°C for 30 minutes, followed by multiple PBS-T washes.

Slides were then processed according to the *Xenium In Situ Gene Expression User Guide* (CG000582 Rev E, 10x Genomics). Gene expression profiling was performed using the Xenium Human Multi-Tissue and Cancer Panel (10x-1000626), targeting 377 genes for prostate, ileum, appendix, pancreas, and colon samples, and the Xenium Human Breast Panel (10x-1000463,) targeting 380 genes for breast cancer samples. Probe hybridization involved probe denaturation at 95°C for 2 minutes, crash-cooling on ice, equilibration to room temperature, and 17-hour hybridization at 50°C. Post-hybridization, slides underwent standardized washes, ligation, amplification, autofluorescence quenching, and nuclear staining following the manufacturer’s protocols.

Slides assembled into Xenium cassettes were processed using the Xenium Analyzer according to the *Xenium Analyzer User Guide* (CG000584 Rev D/E, 10x Genomics). Instrument setup included loading of the Xenium Decoding Consumables Kit (PN-1000487) with appropriate wash buffers and reagents. Slides were scanned to image fluorescently labeled nuclei, and fields of view (FOVs) were selected for sequencing. Data acquisition utilized the Xenium In Situ software suite (v1.9 for standard samples; v2.0 for multimodal segmentation samples), with simultaneous collection of gene expression and segmentation data for multimodal samples.

Post-run, slides were washed with PBS-T and stored at 4°C in the dark until further staining and analysis.

### RNAscope for quality assessment

Chromogenic in situ hybridization was performed on an automated stainer (Leica Bond RX, Leica Biosystems, Deer Park, IL) with RNAscope 2.5 LS Assay Reagent Kit-Red (Cat. # 322150, Advanced Cell Diagnostics, Newark, CA) and Bond Polymer Refine Red Detection (Cat. # DS9390, Leica Biosystems) following manufacturer’s standard protocol. Positive control probe detecting a housekeeping gene (human PPIB, cat # 313908, Advanced Cell Diagnostics) and a negative control probe detecting bacterial (*Bacillus subtilis*) dapB gene (cat #312038, Advanced Cell Diagnostics) were used to assess RNA preservation and detection, and absence of non-specific signal, respectively. The chromogen was fast red and the counterstain hematoxylin. Positive RNA hybridization was identified as discrete, punctate chromogenic red dots under brightfield microscopy. Whole slide images were acquired with an Olympus VS200 slides scanner and VS200 ASW 4.1.1 software (Evident Scientific, Hamburg, Germany) using a 20X/0.80NA objective to generate whole slide images with a pixel size of 0.2738 µm. Image analysis was performed with QuPath 0.5.1 software (Bankhead, P. et al. QuPath: Open source software for digital pathology image analysis ^49^. The positive cell detection and subcellular detection algorithms were used to detect cells and hybridization dots (individual spots and clusters), respectively. The mean number of estimated spots per cell was reported for each sample. The analysis we performed and validated by a board-certified veterinary pathologist (SM).

### Data processing and annotation

For CosMx RNA and protein data, AtoMx Spatial Informatics Platform (SIP) was used for image processing and preprocessing of raw data. A slight modification was applied to the foundational pipeline available on the AtoMx platform (v1.3). For quality control, cells with the following cutoffs were flagged as low-quality cells: total transcript counts less than 20, negative probe counts more than 10%, count distribution 1, area outlier 0.01 (outlier p-value cutoff 0.01). FOV QC was done with default settings (mean, FOV count cutoff 100). The negative control probe quantile cutoff for target QC was set to 0.5, and detection was set to 0.01.

The datasets were normalized by scale factors and total counts, while 6k panels and combined datasets were normalized using total count normalization for scaling. For organ-specific analyses, count matrices were normalized using sc.pp.normalize_total() and log-transformed via sc.pp.log1p() (Scanpy v1.9.3). Dimensionality reduction was performed using PCA, followed by neighborhood graph construction (sc.pp.neighbors()) and UMAP embedding (sc.tl.umap()).

UMAP was generated with the following parameters: 50 PCs for clustering, 40 number of neighbors, 0.01 minimum distance, 5 spread, Cosine distance metric, 0.25 data fraction. Spatial networks were calculated based on the distance (50 um), and neighborhoods based on the Jaccard cutoff of 0.099 and cosine distance. The datasets were exported for cell type annotation and further downstream analysis. For initial automated cell type annotation of RNA data, we used Insitutype ^50^ with organ-specific profile matrices derived from published single-cell and spatial transcriptomics datasets (available via the SAHA GitHub repository; see **Code Availability**). For protein data, cell phenotyping was performed using SCIMAP based on a marker-guided gating strategy. Manual refinement was applied as needed following unsupervised clustering to improve resolution of rare or ambiguous populations. Various batches and organ types were integrated for analysis using Harmony ^51^.

### Organ-specific analyses

To characterize lymphoid architecture in lymph nodes, we applied a multimodal spatial analysis workflow. Spatial niche detection was performed using spaVAE ^52^, a dependency-aware variational autoencoder model. Normalized lymph node datasets were input into spaVAE using default parameters over 50 iterations. The resulting niche clusters were compared to Leiden-based transcriptomic clusters for biological interpretability. Germinal centers (GCs) and peri-GC regions were manually segmented using an interactive workflow implemented with matplotlib.widgets.PolygonSelector and matplotlib.path.Path. Visualizations were generated in Jupyter notebooks using %matplotlib tk, enabling real-time polygon annotation of GC boundaries. Cells were classified based on spatial overlap as intra-GC, peri-GC, or non-GC.

For spatial cell–cell interaction analysis, neighborhood graphs were constructed using Squidpy’s sq.gr.spatial_neighbors() function. Cell–cell interaction matrices were generated with sq.gr.interaction_matrix(), leveraging Leiden-inferred annotations to quantify contact frequencies across annotated cell types within GCs and adjacent regions. Gene expression patterns were visualized using Seaborn’s clustermap() to highlight transcriptional diversity among niche clusters. Spatial autocorrelation statistics were calculated using Moran’s I (sq.gr.spatial_autocorr()), excluding the top 1% of expression outliers. Genes with the highest spatial variability were selected for downstream interpretation and figure display. To identify differentially expressed genes (DEGs) between intra-GC and peri-GC compartments, we applied Wilcoxon rank-sum tests (scipy.stats.ranksums()), followed by multiple hypothesis correction using the Benjamini–Hochberg procedure (statsmodels.stats.multitest.multipletests()). Genes with adjusted p < 0.05 and |logLFC| > 0.5 were considered significant. Volcano plots were generated to highlight key upregulated and downregulated genes across compartments.

### Comparison and integration of CosMx Protein and RNA data

To annotate protein data, we used SCIMAP (v1.3.2), a gating-based hierarchical cell annotation tool which takes a workflow with cell types noted by their marker genes to conduct cell typing (**Extended Data Fig 7a**). Integration of CosMx RNA and protein data was performed using MaxFuse (version 0.02). a Python (version 3.8.20) package that uses co-embedding, data smoothing, and cell matching to integrate “weakly-linked” multi-modal omics data that have little overlapping features ^10^. Broad and granular annotations from RNA samples are integrated into serial sectioned protein samples to generate protein cell type annotations. We generated cell type mapping to analogous RNA and protein regions to create visual comparisons. To conduct the concordance comparison between SCIMAP and Maxfuse annotations, we examined how cells called by one modality are called by the other by calculating adjusted rand index and mapping a matrix of cell counts for the cell types of both modalities as a heatmap.

For RNA-protein correlations and concordance comparisons, all analyses were conducted using R version 4.4.3 and the following R packages: Seurat (v5.2.1), reticulate (v1.41.0), anndata (v0.7.5.6), dplyr (v1.1.4), ggplot2 (v3.5.2), ggrepel (v0.9.6), reshape2 (v1.4.4), and RColorBrewer (v1.1.3). No cell-wise or feature-wise filtering was applied to either the transcriptomic or proteomic datasets for this analysis. We identified 53 features common to both RNA and protein modalities across 15 matched tissue cross-sections from 9 different organs. For each tissue pair, the mean RNA and protein expression levels for each common feature were computed after excluding values equal to zero, to avoid bias from low-input measurements. Mean values were also calculated at the organ level by averaging feature expression across all tissue sections from the same organ. In addition, pooled averages were computed by aggregating all features across all organs. To assess the relationship between RNA and protein expression, we computed the log10-transformed mean expression values of each feature and visualized them using scatter plots, with log10(RNA mean) on the y-axis and log10(protein mean) on the x-axis. These plots were generated for tissue pairs, organs, and the aggregated dataset. Linear regression models were fit to each of these comparisons using the lm() function in base R, with the following formula: log10(RNA.mean) ∼ log10(PRT.mean). We also generated analogous plots and regressions to compare average RNA and protein expression values across all common features within each tissue pair and organ. Spearman’s rank correlation coefficient was used to calculate RNA-protein correlations across organs and genes.

### Comparative analysis of H&E images with spatial transcriptomics datasets

WholeLslide H&E imaging data was processed using LazySlide ^53^. Gastrointestinal tract samples (appendix, colon, ileum) were segmented and tesselated with tiles of 512 pixels at 0.5 µm/pixel (∼20X magnification). For each slide, we randomly sampled 250 tiles and extracted highLdimensional morphological embeddings leveraging the CONCH multiLmodal vision– language model ^15^. Subsampling was performed to ensure equal contribution of various slides/tissue types. Tile embedding values were scaled, reduced with Principal Component Analysis (PCA) and visualized using the Isomap (scikit-learn) or Uniform Manifold Approximation and Projection (UMAP) algorithm ^54^ and clustered using the Leiden algorithm ^55^ at resolution 0.5 with Scanpy ^56^. For downstream interpretation, we assembled a panel of 79 histolopathological terms related to instestinal biology or pathology. We computed the similarity between the text embedding of each term in each CONCH feature vector of each tile. Text embedding term values were compared between morphological clusters using a Wilcoxon test, and the top 3 most differential terms were used to describe each cluster.

In parallel, we applied CellViT ^57^ to the same wholeLslide images to perform endLtoLend nuclear segmentation and classification. Within each sampled tile, we quantified cellLtype abundances (e.g., epithelial, immune, connective) and averaged these cellular proportions per morphological cluster previously derived by the CONCH features. To link tileLbased morphological phenotypes with molecular cellLtype composition, we performed canonical correlation analysis (CCA) implemented in scikit-learn between the relative abundance of each morphology cluster and the corresponding cellLtype proportions inferred by the CosMx platform per tissue slide.

### Integrative analysis of SAHA COL and CRC CosMx data

Downstream analysis was performed using Seurat (v5.0.1) ^43–47^ throughout to assess cell types and perform graph-based clustering. To harmonize gene probes across both datasets, gene probes targeting closely related genes (previously combined in an earlier spatial probe set but separated into individual genes in the current set) were merged. Specifically, genes originally represented as single combined probes (e.g., *CCL3/L1/L3, CXCL1/2/3*) were merged into single meta-probes by summing their expression values across individual gene measurements. A complete list of merged genes is available in **Supplementary Table 3**. Genes without common representation across both spatial datasets were excluded to ensure consistent comparisons. After matching the probes, each FOV in the CRC sample was matched to an FOV from the COL sample to achieve relatively equal cell numbers (Figure 7, Extended Data Fig. 8).

Following this process, raw count matrices from each object were exported and combined using the merge() function from Seurat. Normalization, scaling, and PCA calculation were performed using standard Seurat functions. Objects were then integrated using Seurat’s IntegrateLayers() function with the Harmony algorithm ^51^ (parameters: theta = 2, lambda = 0.1). After integration, Uniform Manifold Approximation and Projection (UMAP) embedding was performed using 30 principal components (PCs). Clusters were defined using a resolution of 0.3, and low-quality cells (Cluster 11) were excluded based on low feature counts. Prior to marker identification, the JoinLayers() function was applied as recommended by Seurat developers. Cluster markers were identified using Seurat’s FindAllMarkers() function (parameters: min.pct = 0.2, logfc.threshold = 0.5, adjusted p-value < 0.05; Supplementary Table X).

Differentially expressed unique genes highlighted in Fig. 7c were identified with the FindMarkers() function using the MAST test (min.pct = 0.1, logfc.threshold = 0.5, adjusted p-value < 0.05; Supplementary Table X). Spatially differentially expressed genes (Fig. 7d-f) were computed using the Voyager tool ^58^ to calculate Moran’s I. Field of views (FOVs) were clustered based on the top 200 spatially variable genes per field using principal component analysis (25 PCs), followed by k-means clustering (k = 2) as visualized in Fig. 7d.

Visualizations were generated with Seurat’s functions: UMAPs (Fig 7a, Extended Data Fig. f,h) (DimPlot), Dotplots (Fig. 7h-2, Extended Data Fig. b,g) (DotPlot), violin plots (Fig 7c and e, Extended Data Fig. a) (VlnPlot), and pseudo-spatial images (Fig 7g-4, h-2, Extended Data Fig. h-1) (ImageDimPlot). Bar plots (Fig 7g-3, Extended Data Fig. c,f) were created using dittoBarPlot from dittoSeq ^59^, and spatial crypt visualizations were generated using ggplot2’s geom_point() (Fig. 7b,f, Extended Data Fig. d,e). Cell-cell interaction analysis (Fig. 7h-3) was performed using CellChat software ^60^ (conversion factor = 0.18; interaction range = 250; scale.distance = 4.5; contact range = 20).

### Data explorer generation

All generated data were made available for interactive exploration using Vitessce ^61^. For each individual sample, cell segmentation polygons were converted into a JSON format in which each cell is represented as an object containing a list of [x, y] coordinate pairs corresponding to its boundary vertices. The visualization interface included a “Spatial” viewer, which displays the sample in physical (x, y) space, allowing users to explore gene expression for any gene alongside cell-level annotations derived from downstream analyses. Additional panels included a “Quality Control” window showing violin plots for metrics such as transcripts per cell and unique genes per cell, a “Features” window listing available genes, and a “Cell Labels” window summarizing cell annotations. All panels were linked, enabling dynamic cross-filtering: selections made in one viewer were automatically reflected across the others. Files generated for the Vitessce viewers are hosted on Zenodo and the script used to generate these files was deposited on github (See **Data and Code Availability**).

### Data and Code Availability

Raw sequences, images, and processed datasets from single-nuclei sequencing and spatial transcriptomics are available via Zenodo (https://zenodo.org/communities/saha) and web portal (https://saha-project.org), and code blocks used to analyze the data can be accessed in the GitHub repository (available upon acceptance; https://github.com/saha-project).

## Acknowledgments

We thank the WorldQuant, GI Research Foundation, Bumrungrad International Hospital, the National Institutes of Health (R01MH117406), and the LLS (MCL 7001-18, LLS 9238-16, 7029- 23). E.A., Y.Z., and A.F.R received funding from Angelini Ventures S.p.A. Rome, Italy. P.S.D. thanks NIDDK (1U01DK134321, 5R01DK135620), NIAID (1U19AI181102), and Digestive Health Foundation for the support. We would like to acknowledge former SAHA members from NanoString Technologies, whose contribution was crucial for establishing and generating the early phase of the project, specifically Nathan Schurman, Joachim Schmid, Raymond Tecotzky, Sarah Church, and Charlie Glaser. Most importantly, we thank the patients and their family members who provided valuable samples for this research.

## Author contributions

Conceptualization: J.P., C.E.M., R.E.S., S.L.H., A.A., R.D.G., E.H.

Patient sample coordination: E.H., B.R., R.C., S.P., F.S.

Experiment and data generation: A.A., R.D.G., H.C., S.M., P.S.D., Y.L., L.P., K.Y., A.H., M.K., D.M., L.W., A.W., M.P., A.W., Y.L.

Investigation and data visualization: J.P., E.O., N.B., F.S.D., L.Z., M.M., Y.Z., E.A., J.K., M.R.A., E.M., J.R., Jac.P., A.A.A., P.D.,

Data management and portal development: J.P., T.N., M.M., F.S.D., Jac.P. Supervision: C.E.M., R.E.S., S.L.H. A.F.R., J.T.P., L.M., O.E., G.C., M.L., J.B., Ja.K.

Writing: J.P., R.D.G., F.S.D., L.Z., Y.Z., A.F.R., E.A., A.A.A., E.O., N.B., J.T.P., C.E.M., S.B., S.R., P.D., S.M.L., A.B., T.N., C.E.M.

All authors discussed the results and contributed to the final manuscript.

## SUPPLEMENTARY TABLES

Supplementary Table 1. SAHA Cohort Demographics and Clinical Metadata.

Summary of demographic and clinical characteristics of the SAHA cohort, including patient age, sex, tissue of origin, comorbidity (if applicable), and sample designation.

Supplementary Table 2. SAHA Experimental Run Metadata.

Detailed metadata for all SAHA spatial transcriptomics runs, including information such as slide/sample identifiers, tissue types, assay panels, staining protocols, quality metrics/statistics, platform used (e.g., CosMx, Xenium), and batch/run identifiers.

Supplementary Table 3. Additional Information related to CRC Analysis.

Additional files regarding the CRC and COL comparative analysis including list of probe substitutes, differential gene expression lists, cell marker lists, full result list of Moran’s I results.

---

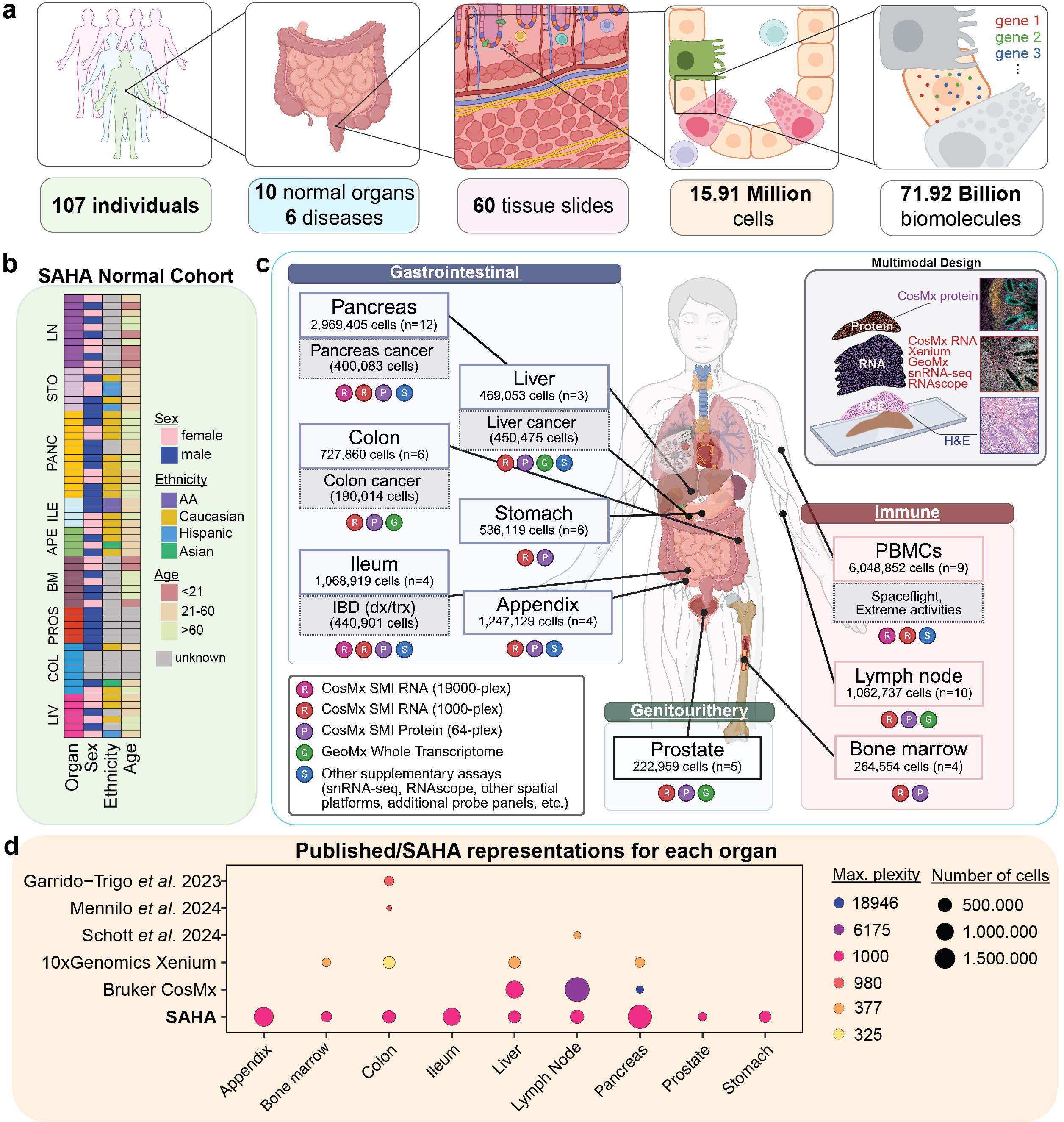

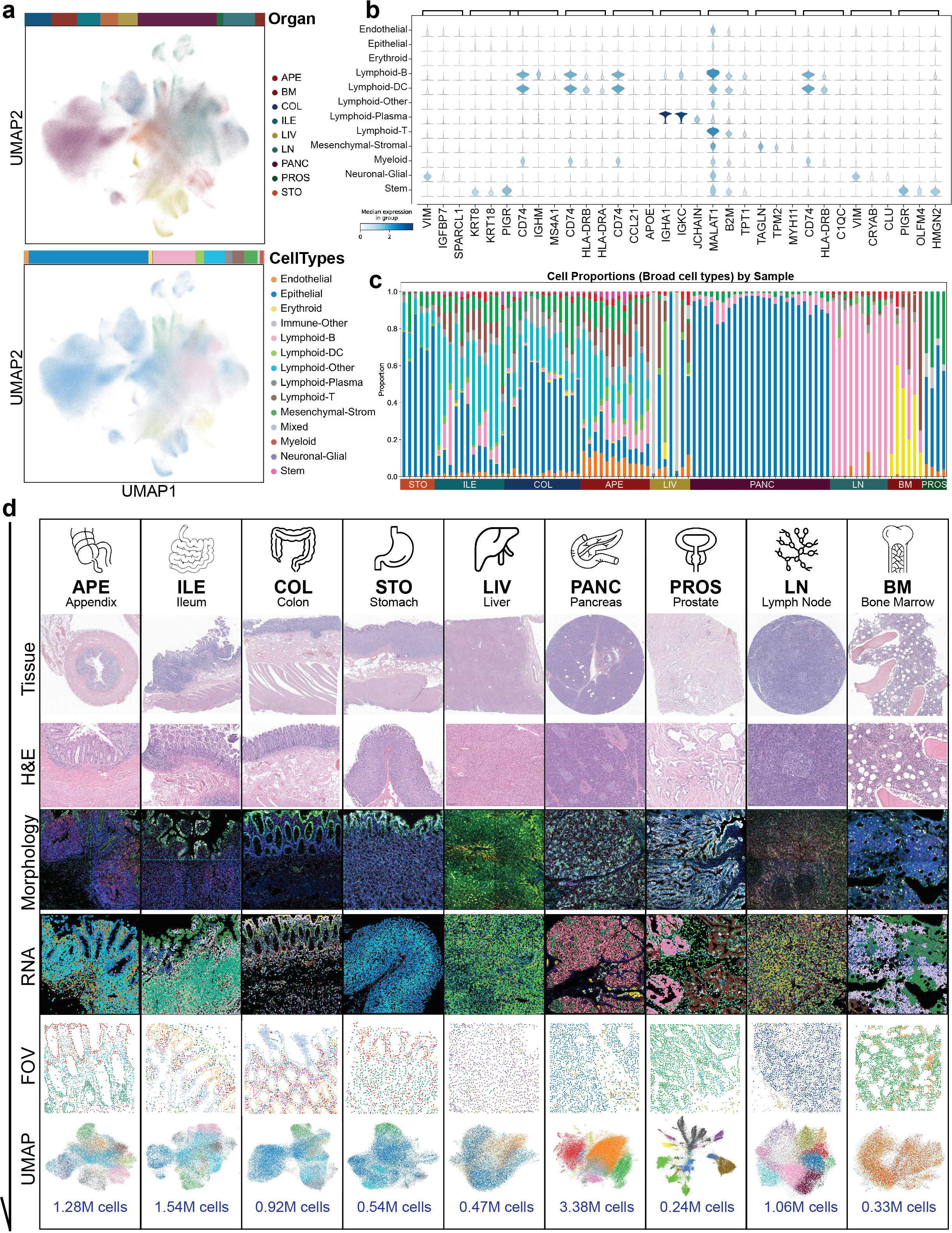

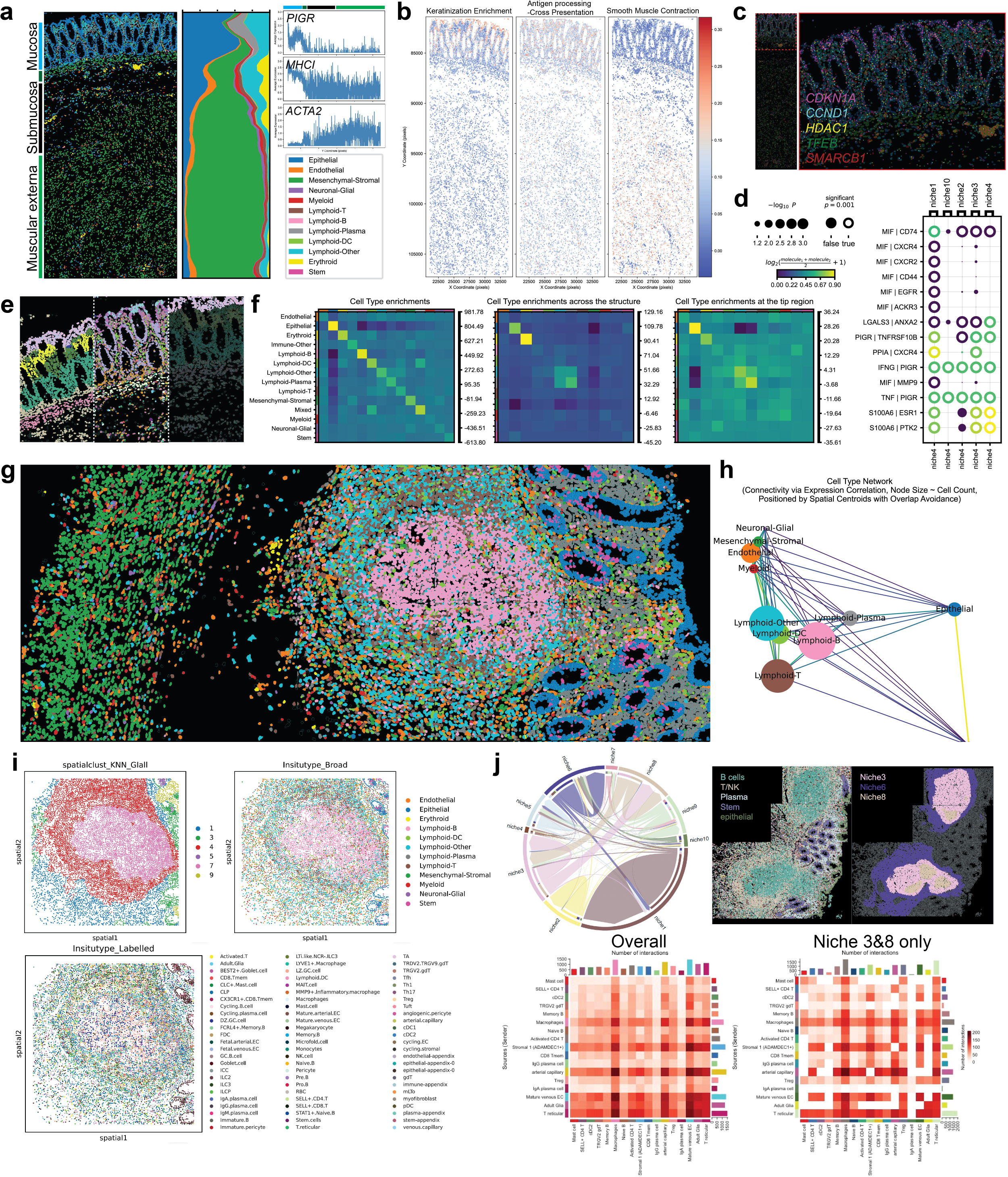

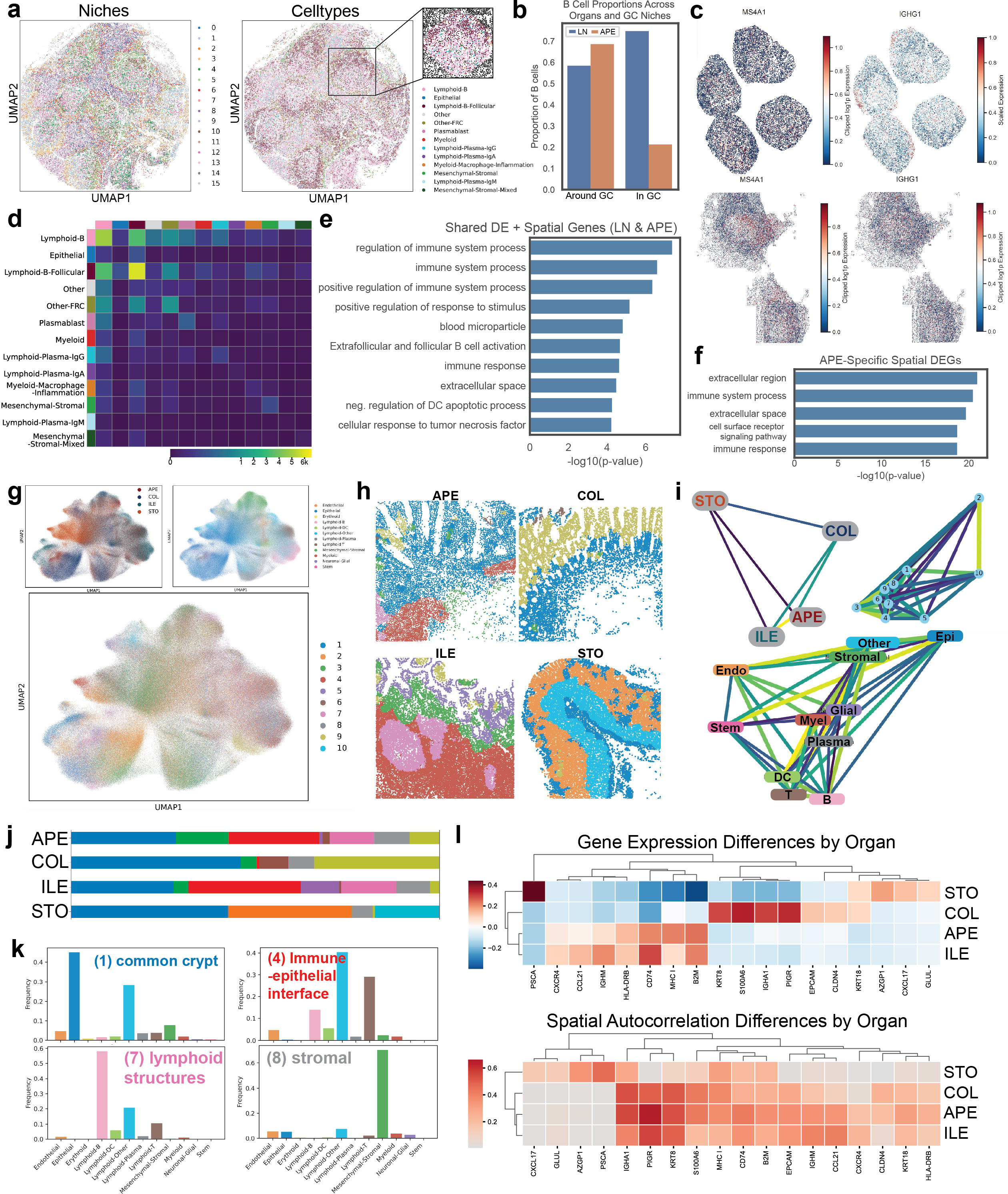

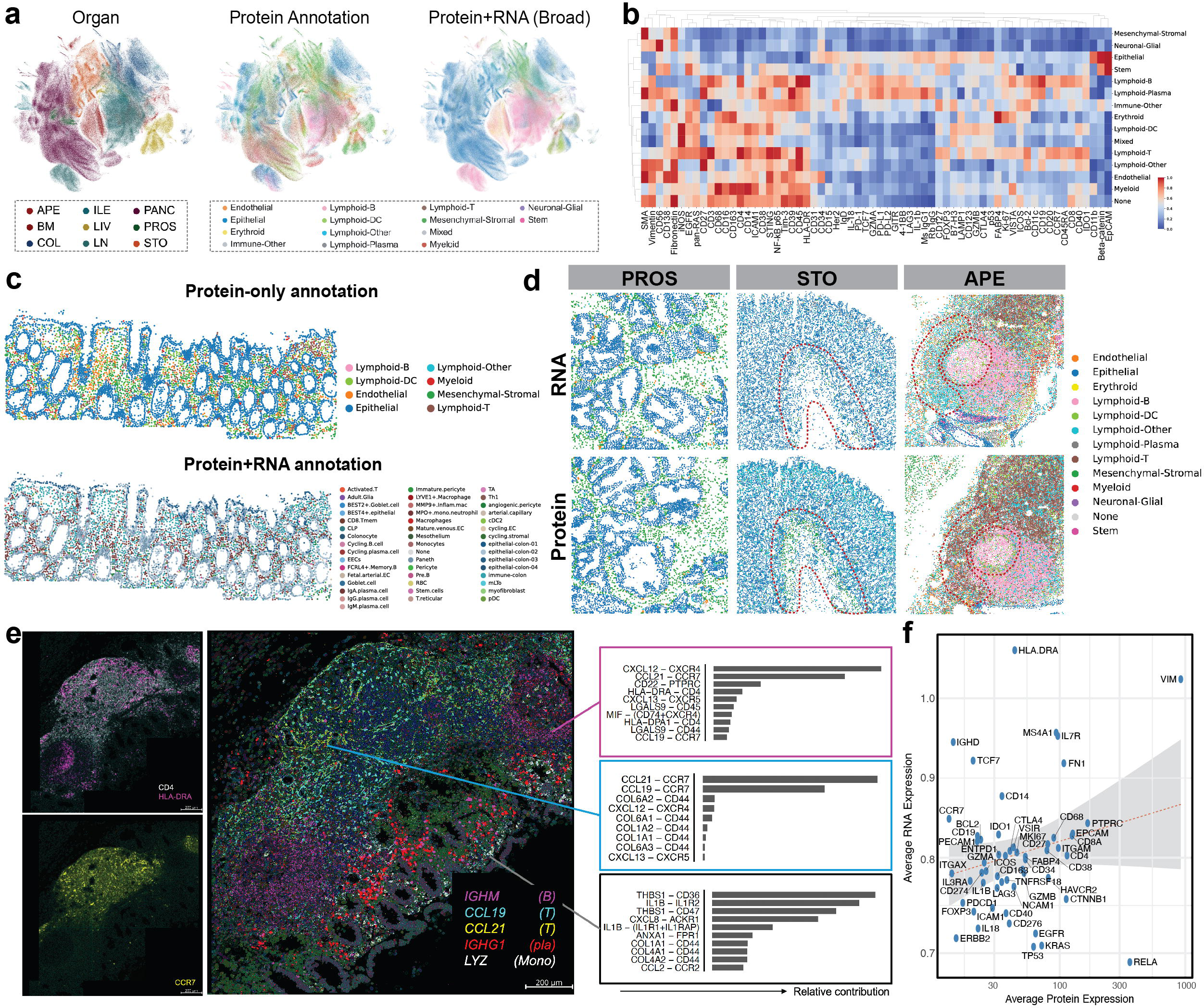

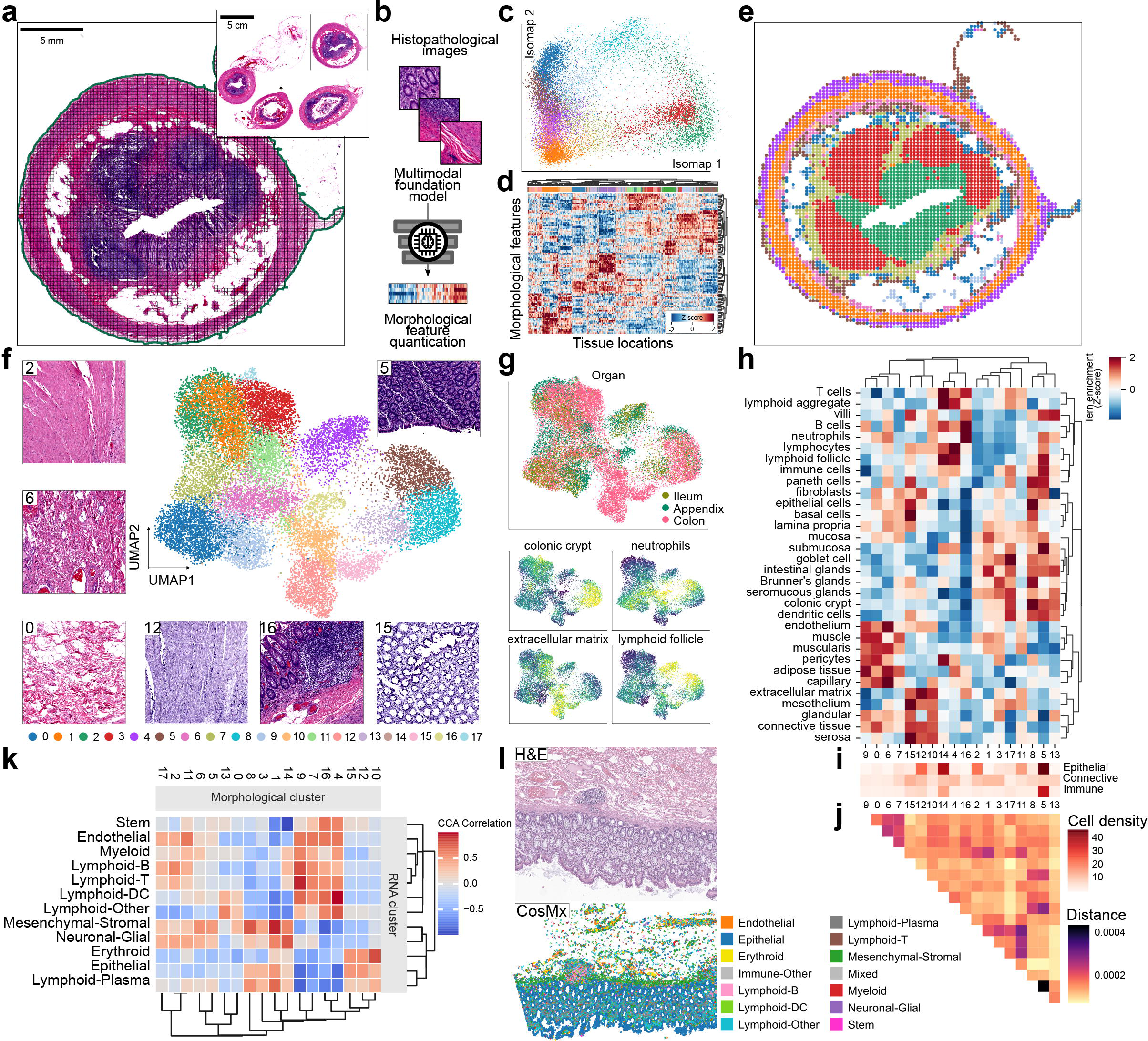

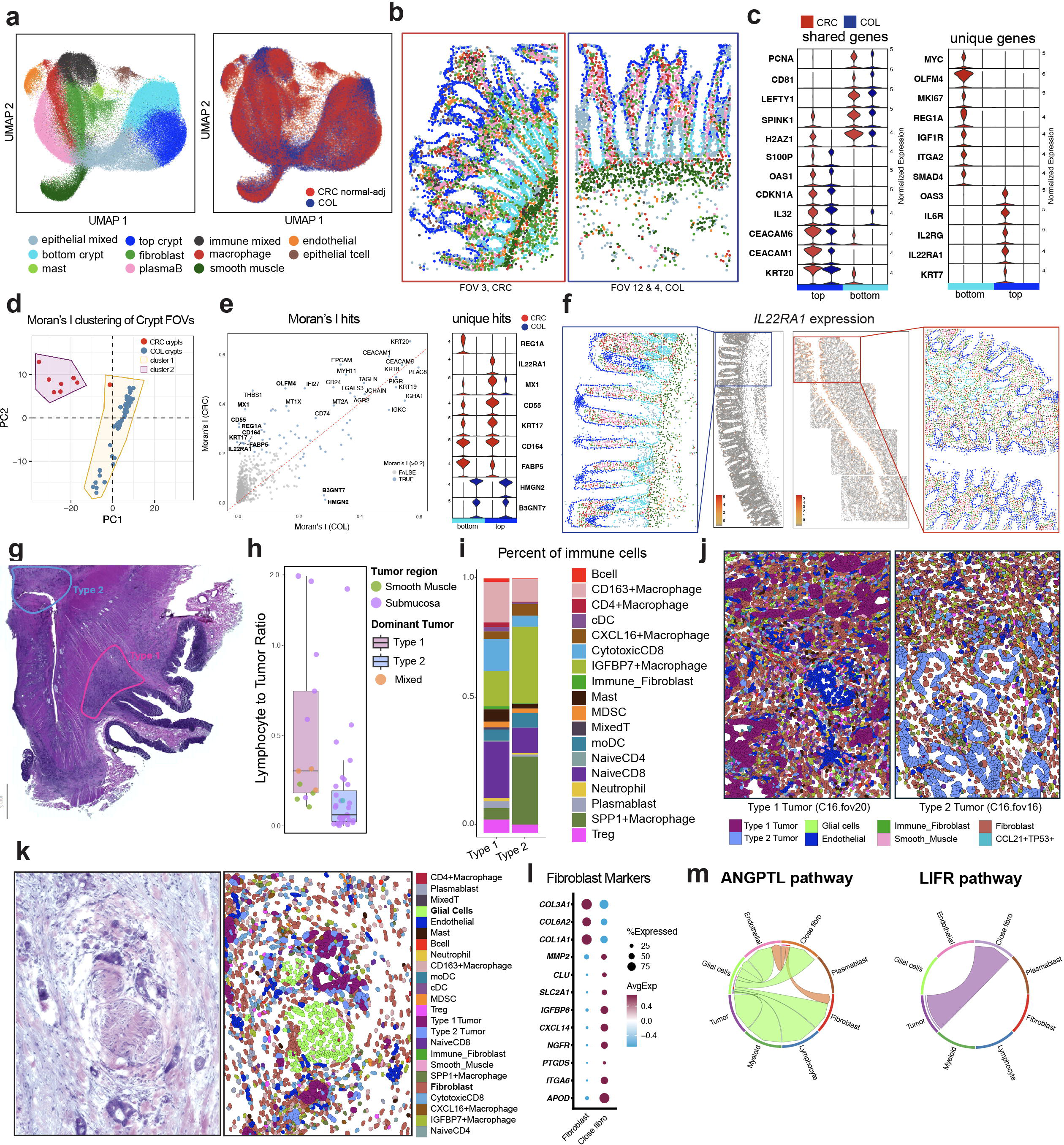

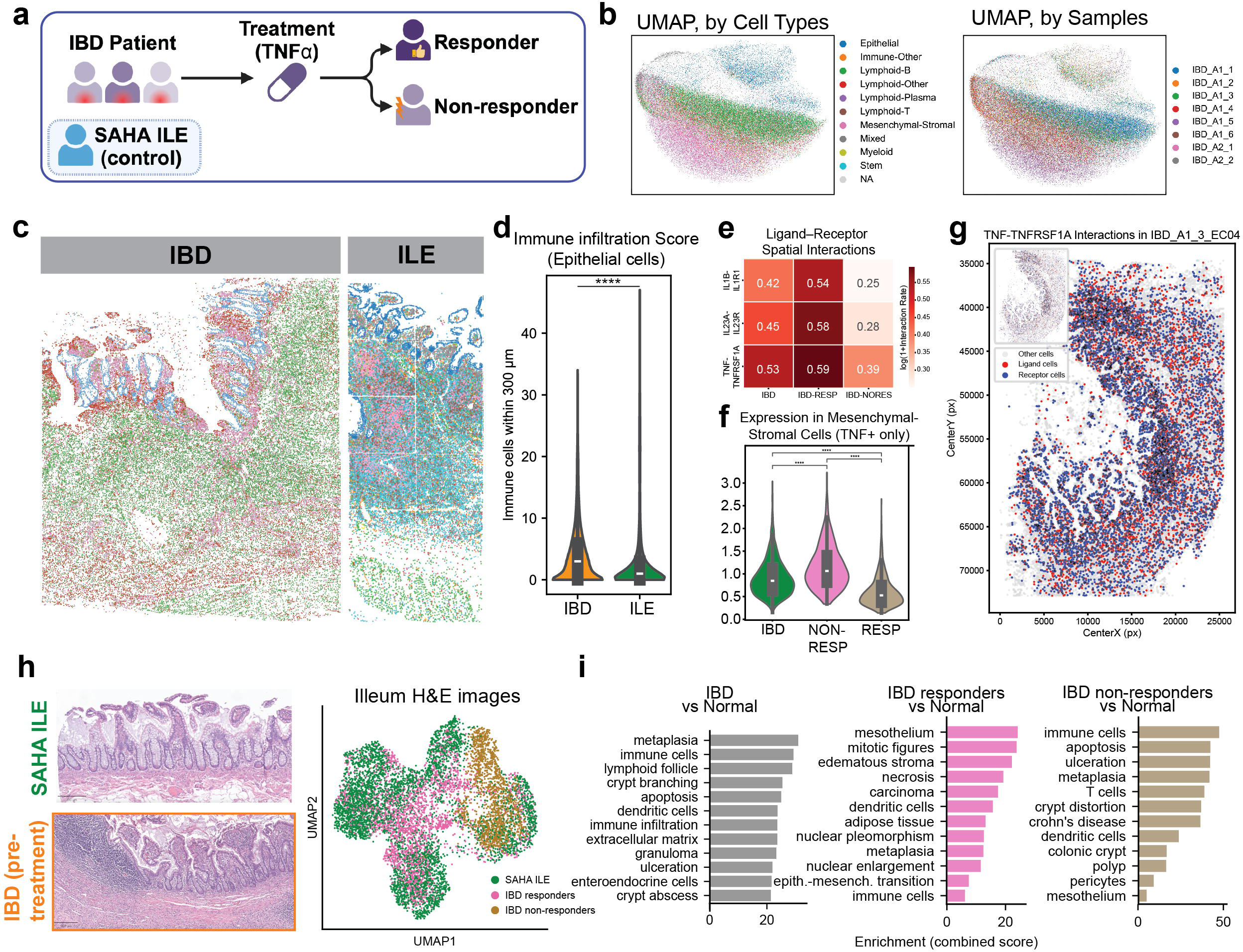

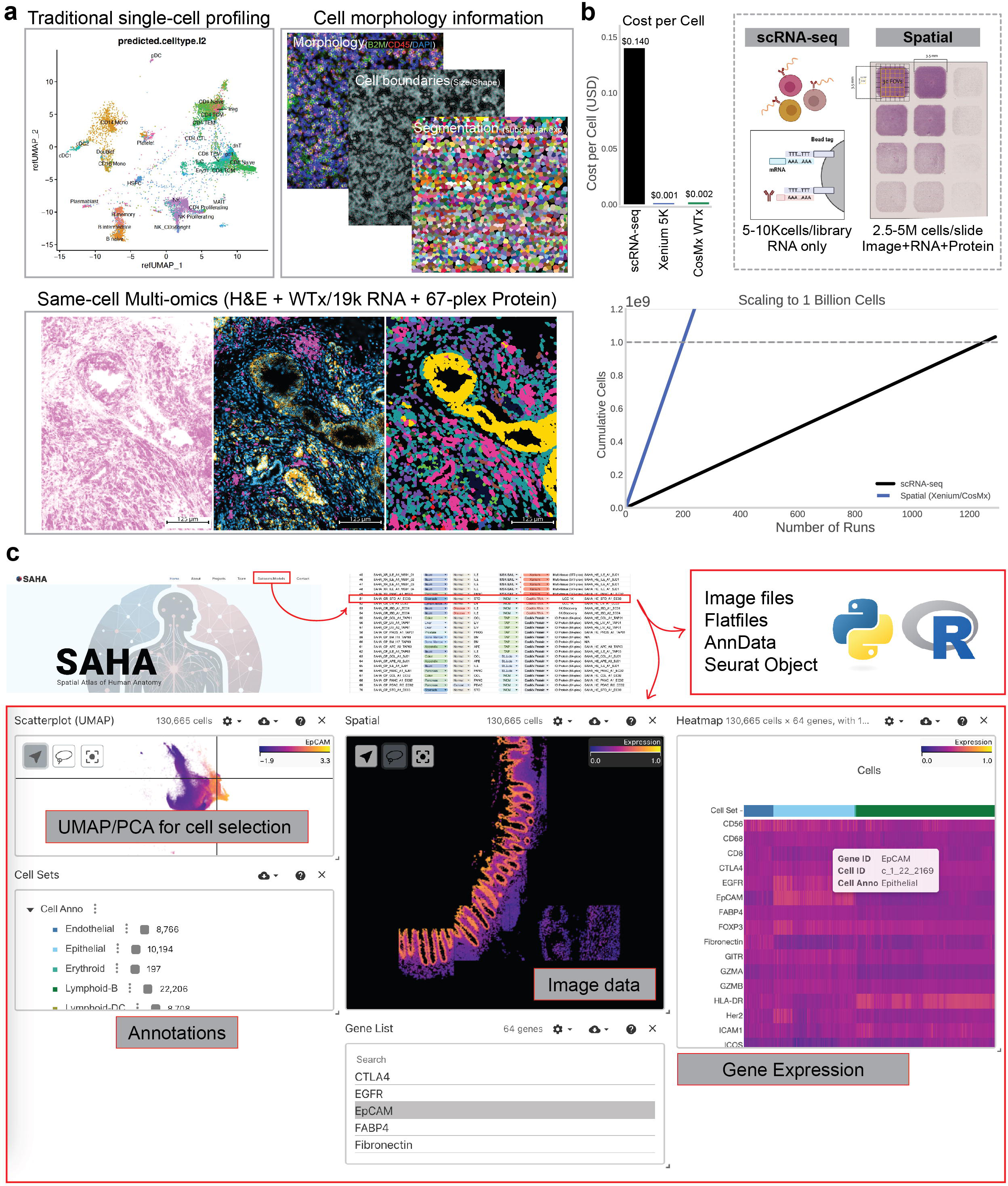

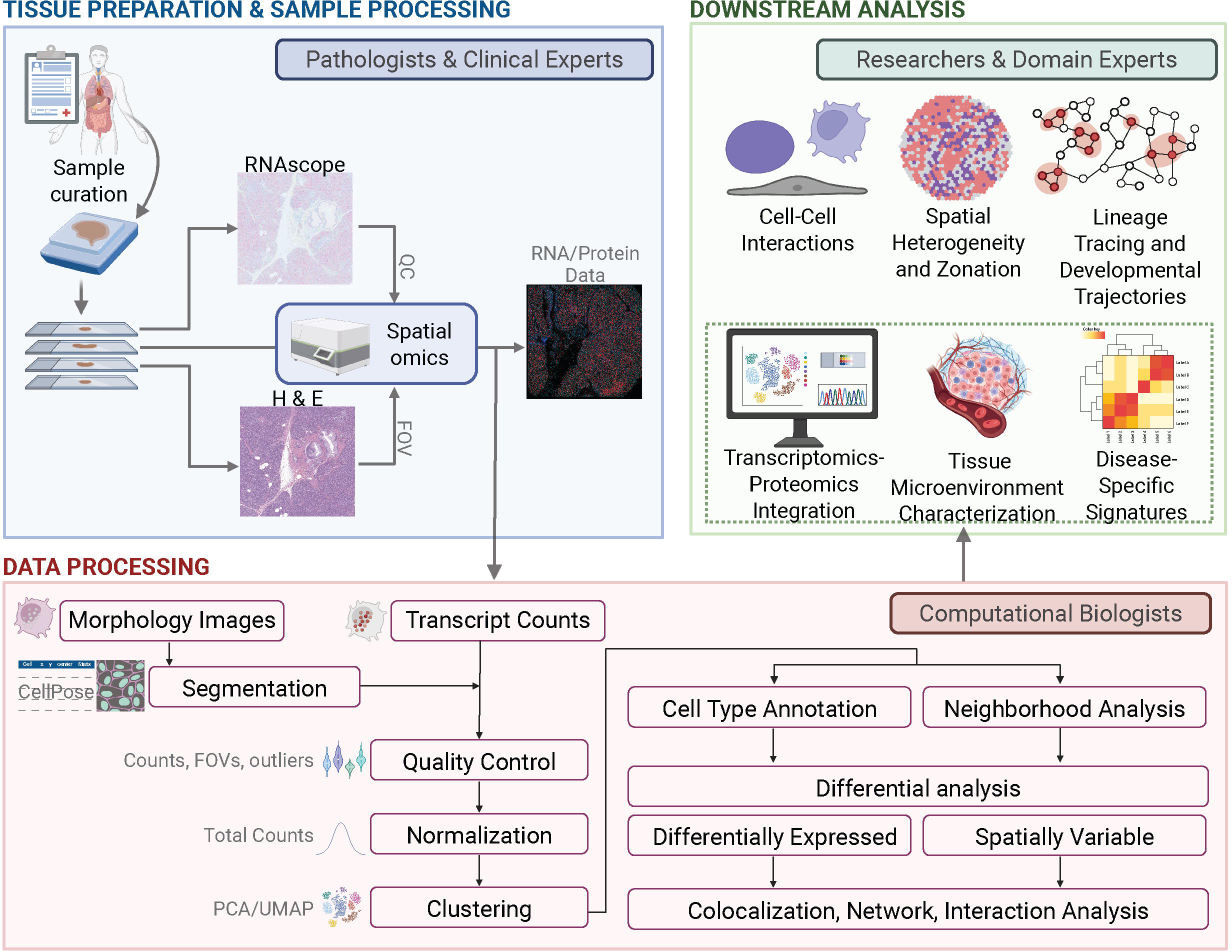

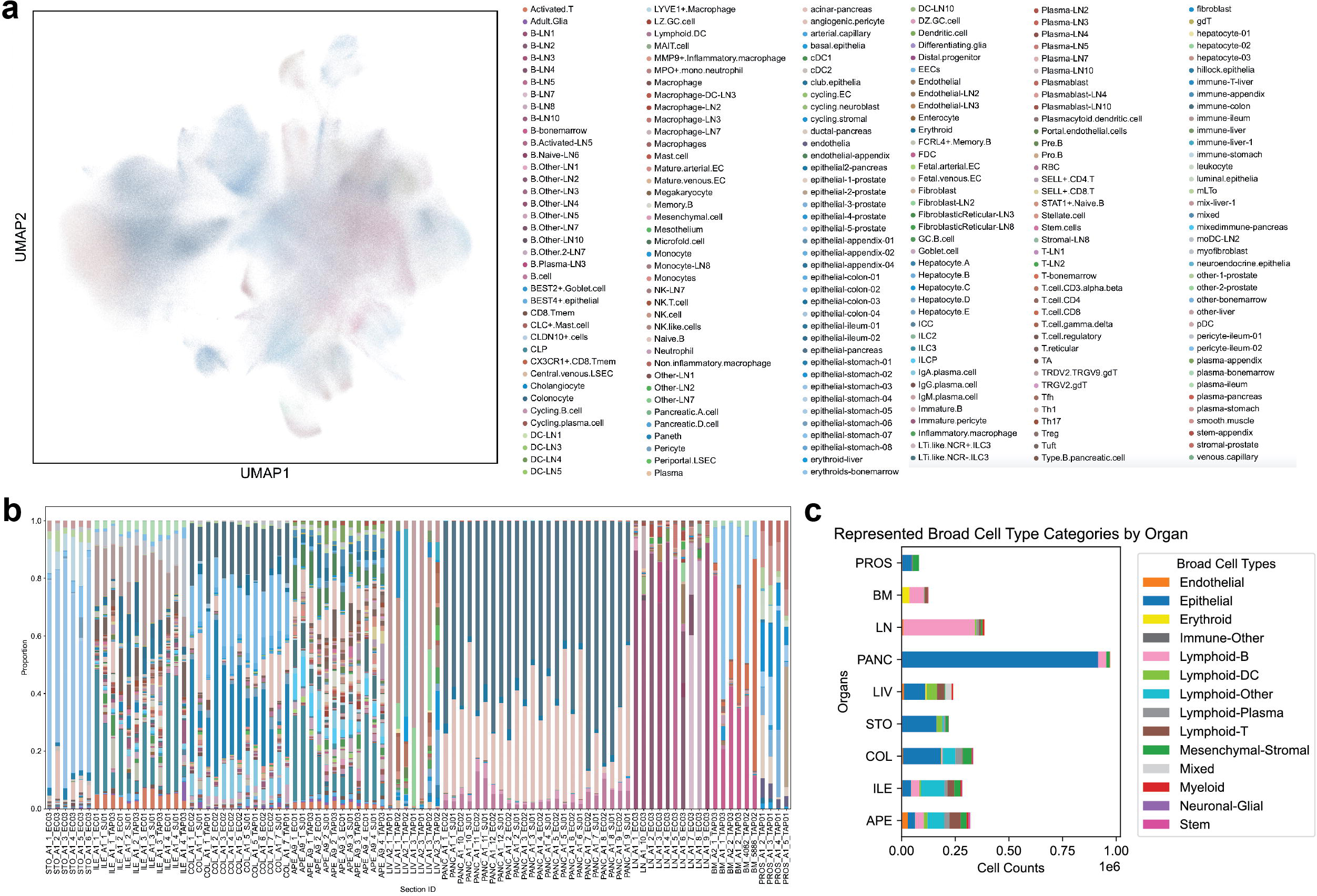

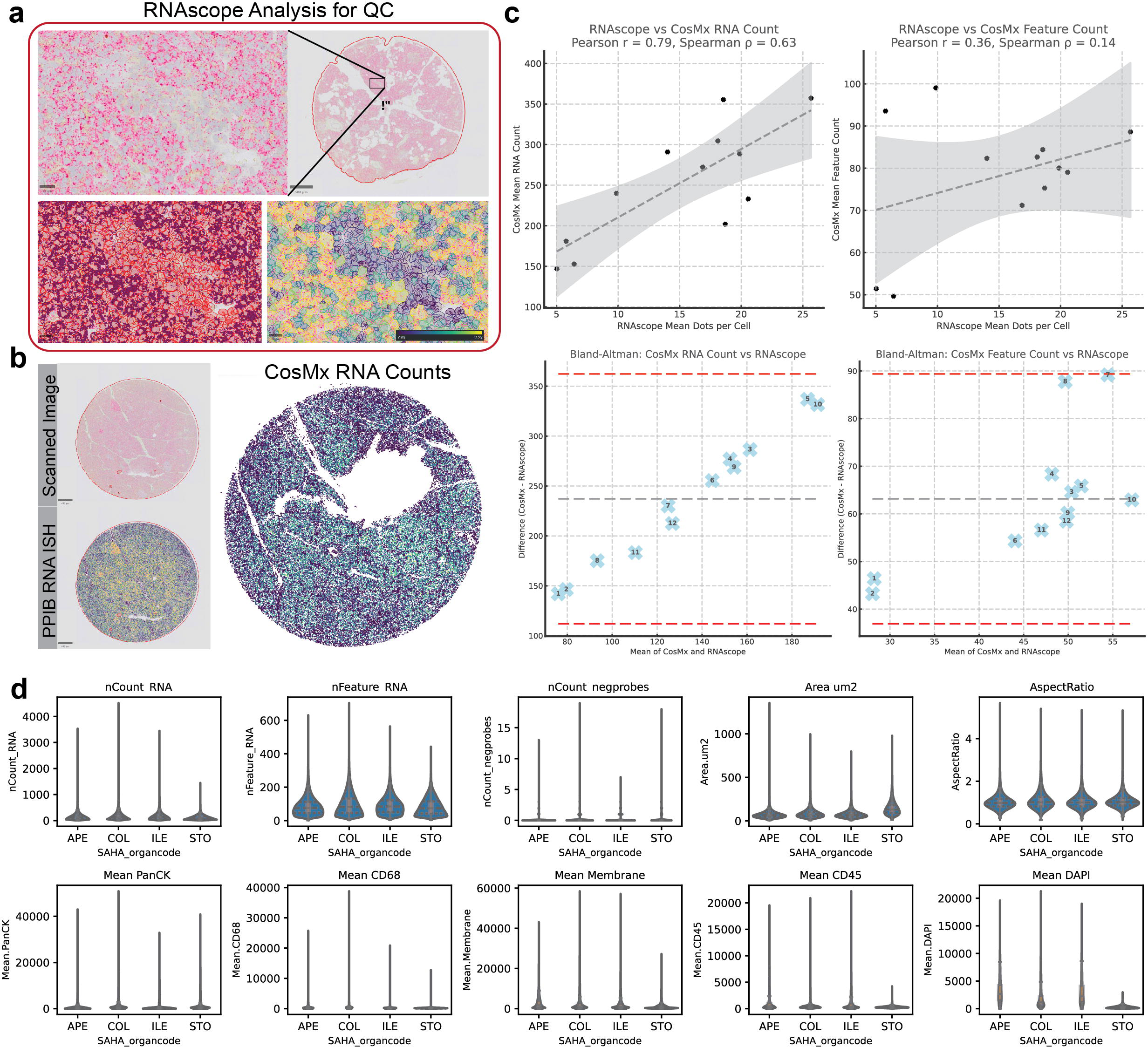

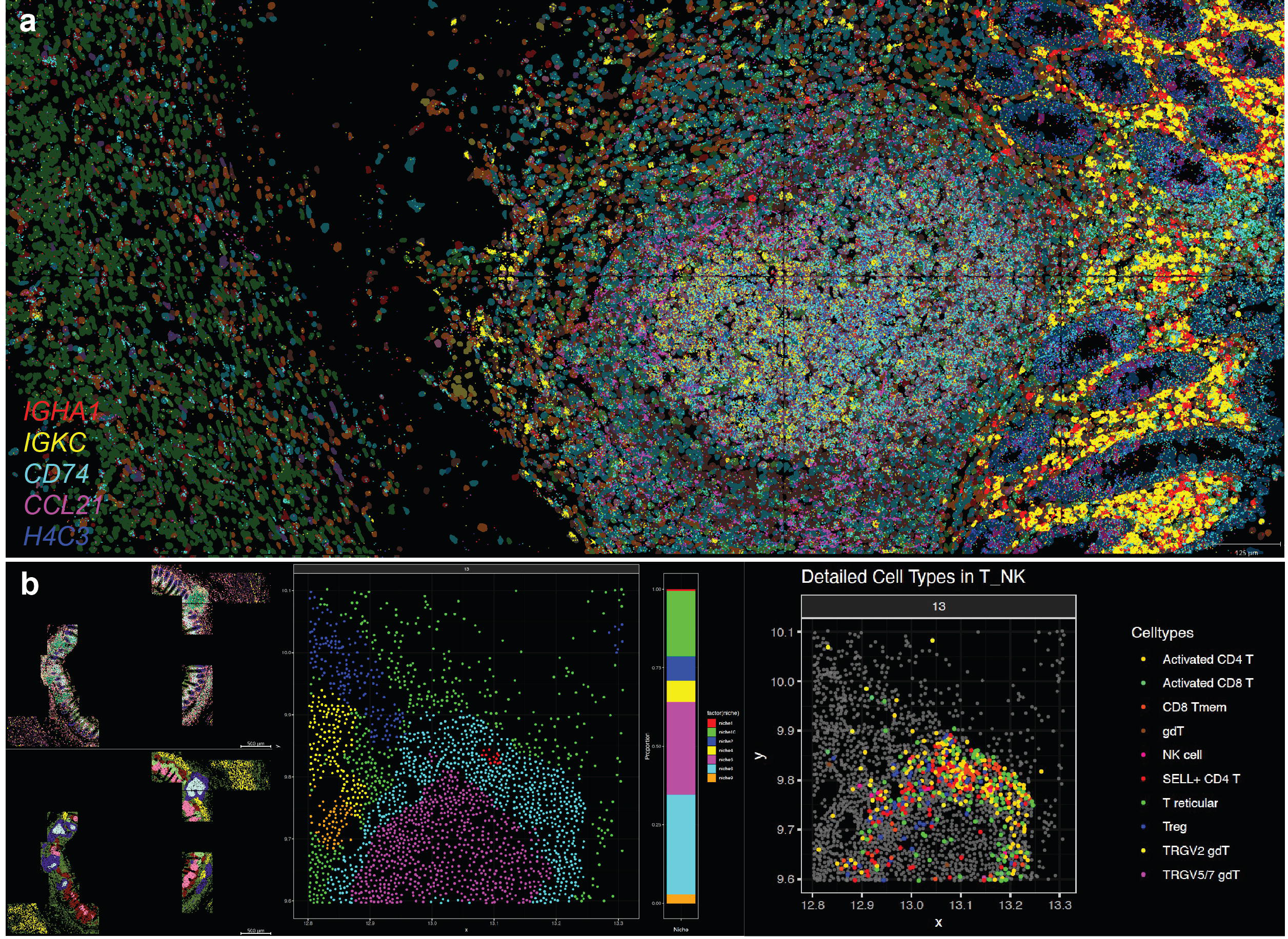

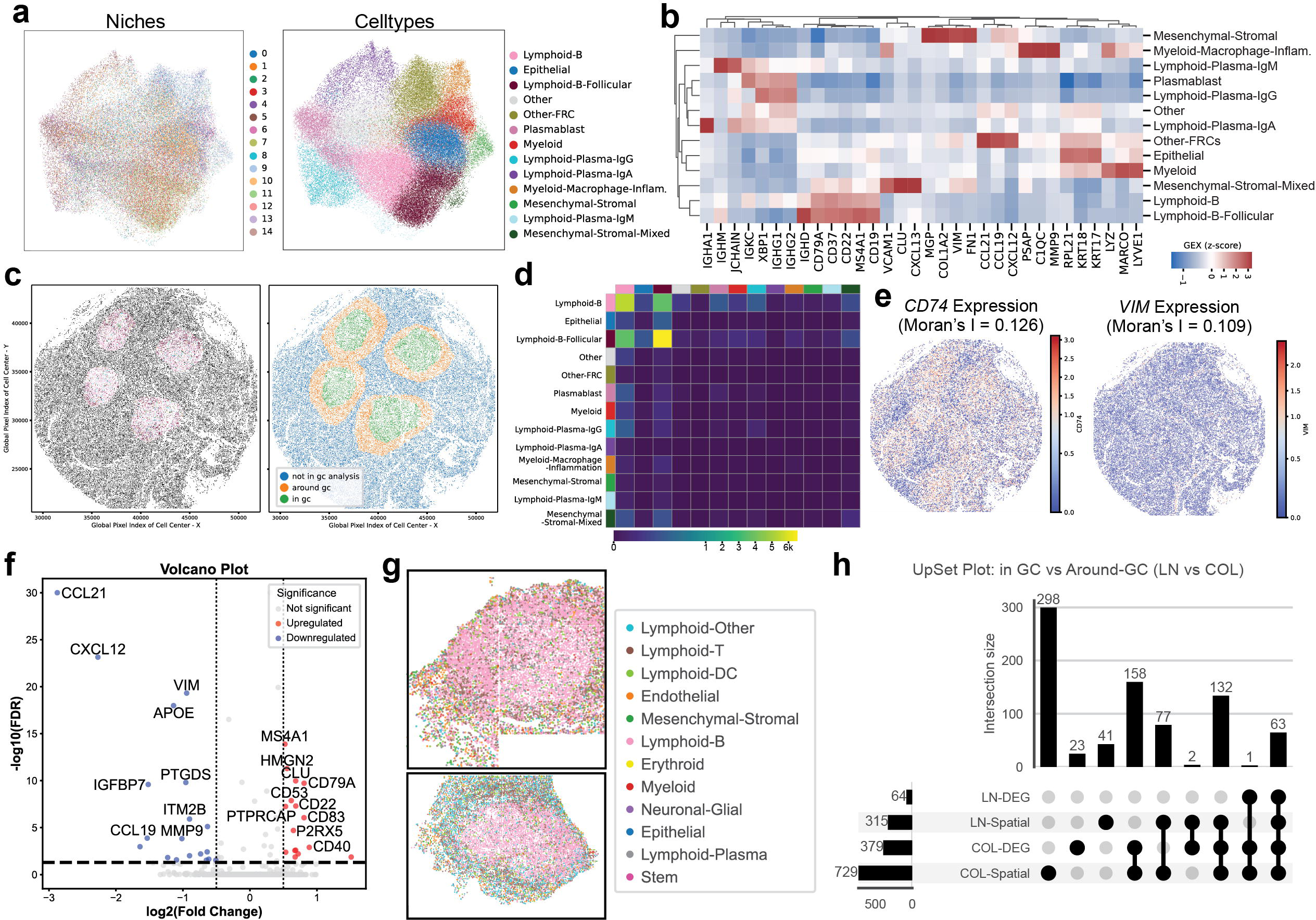

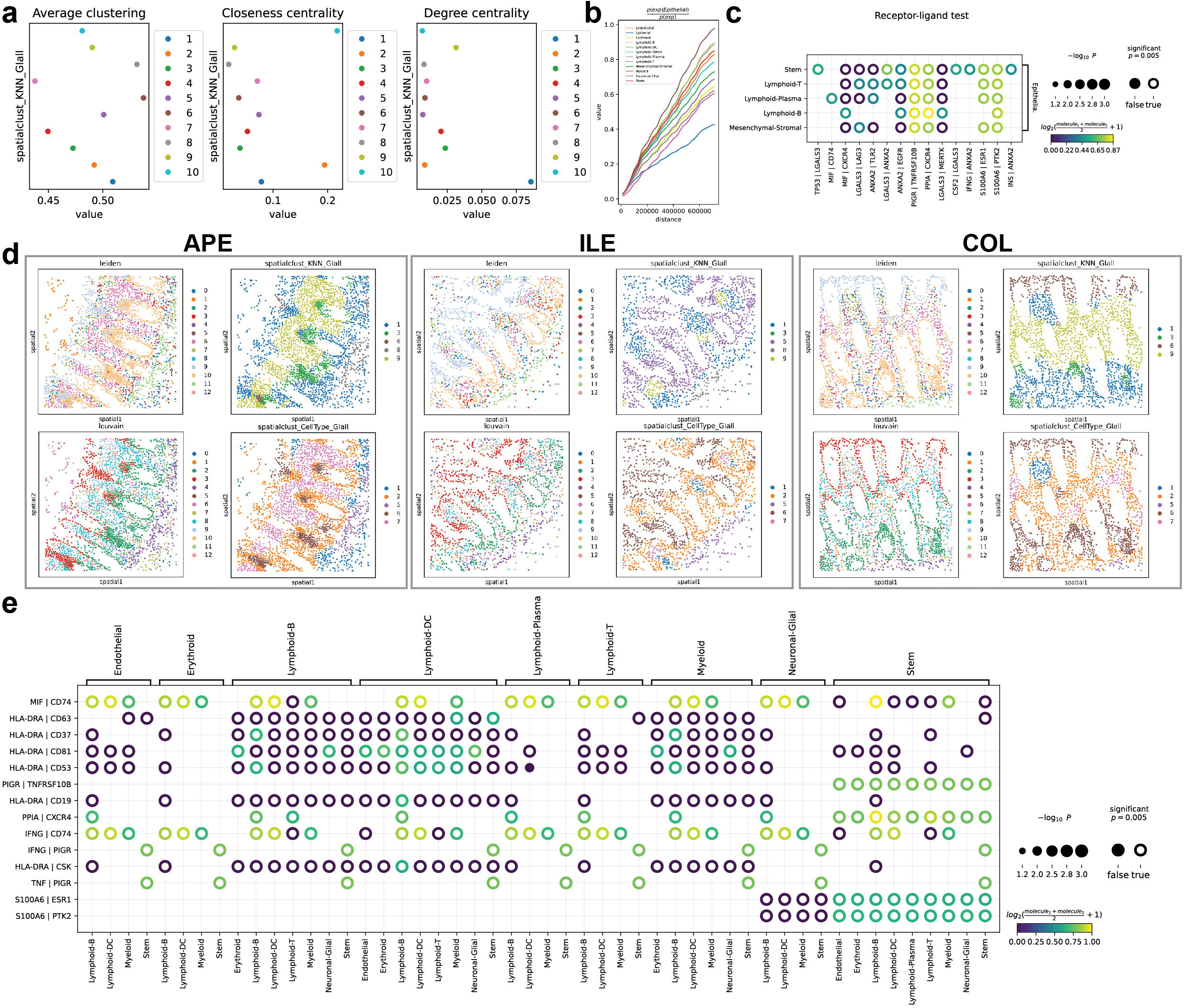

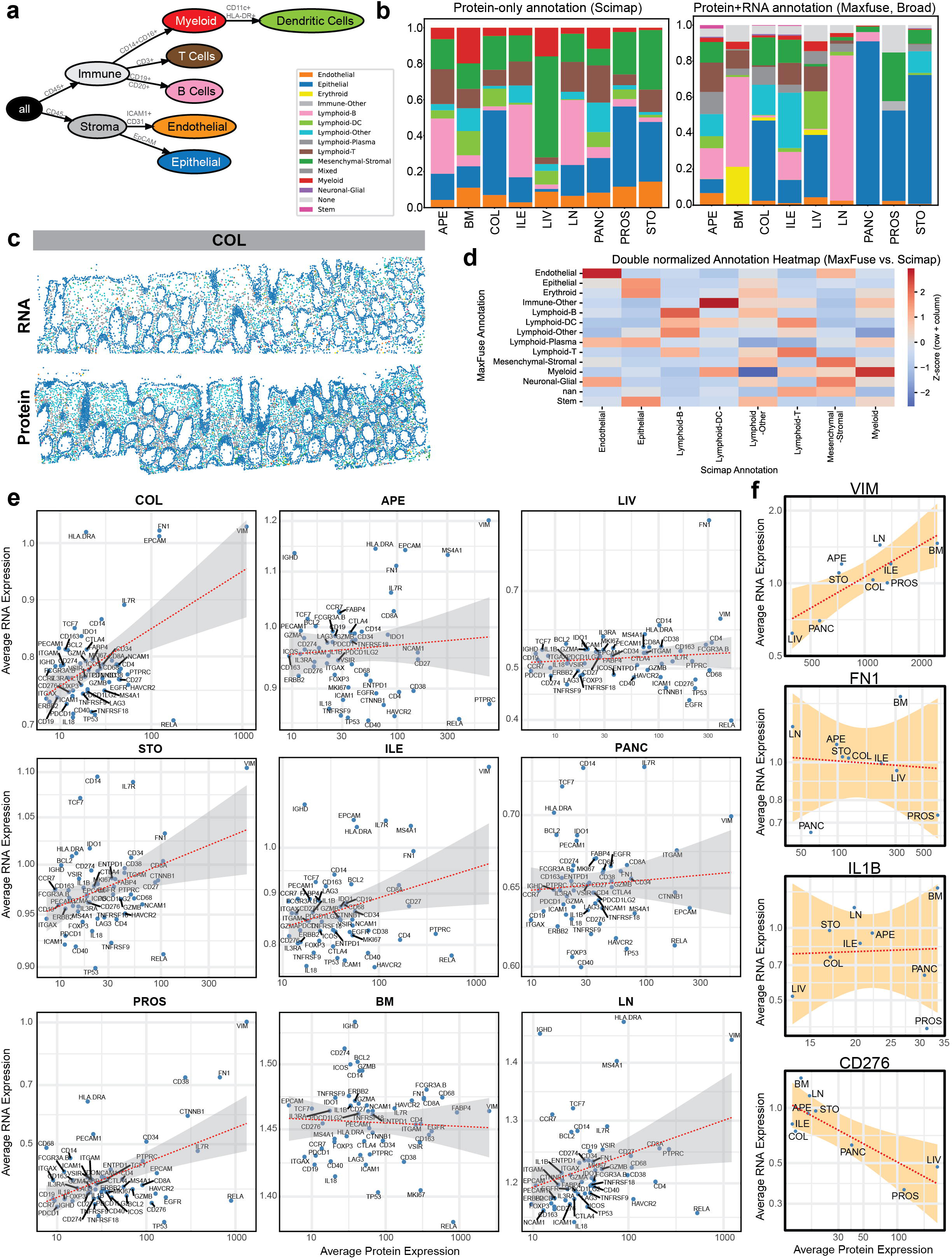

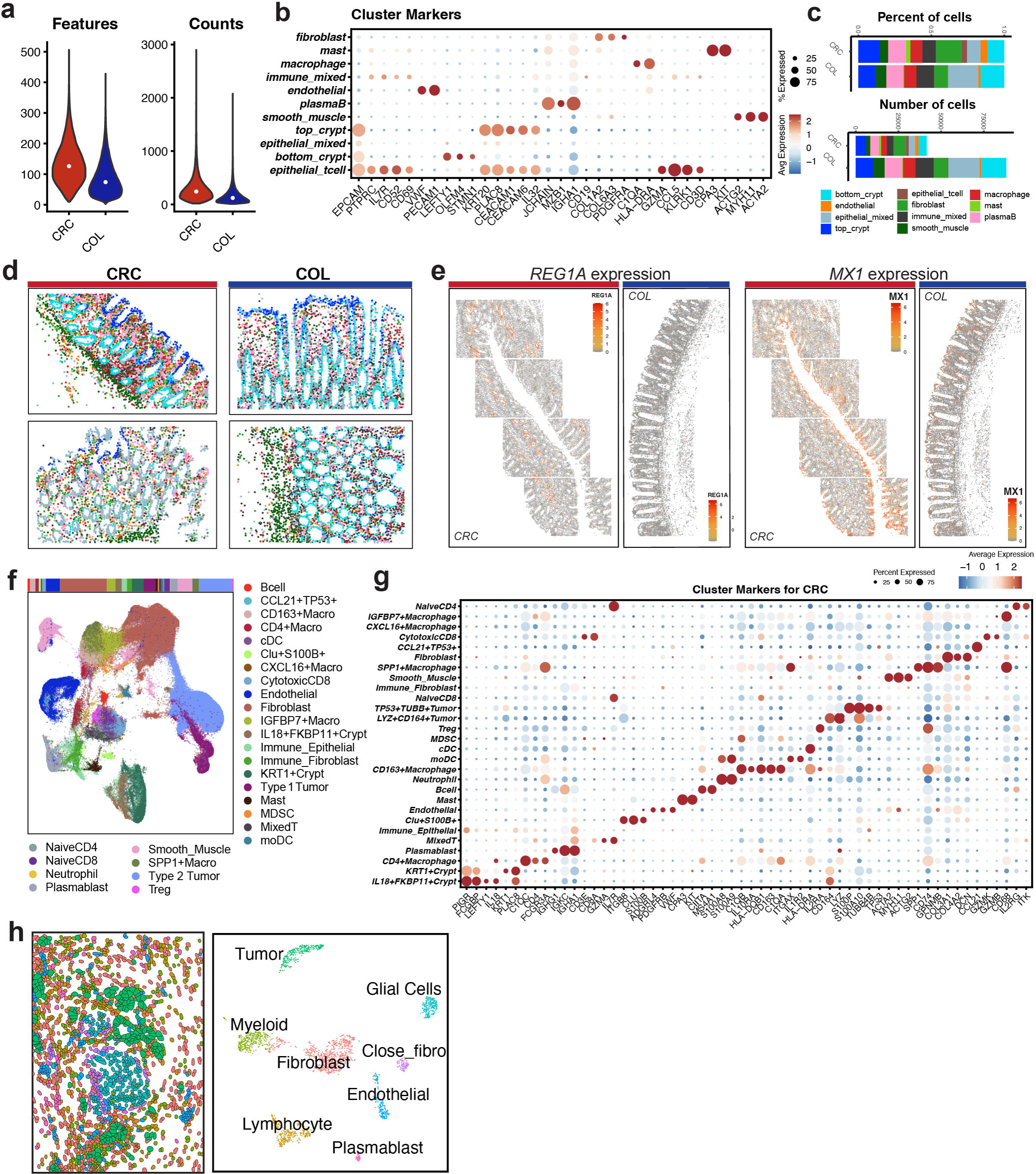

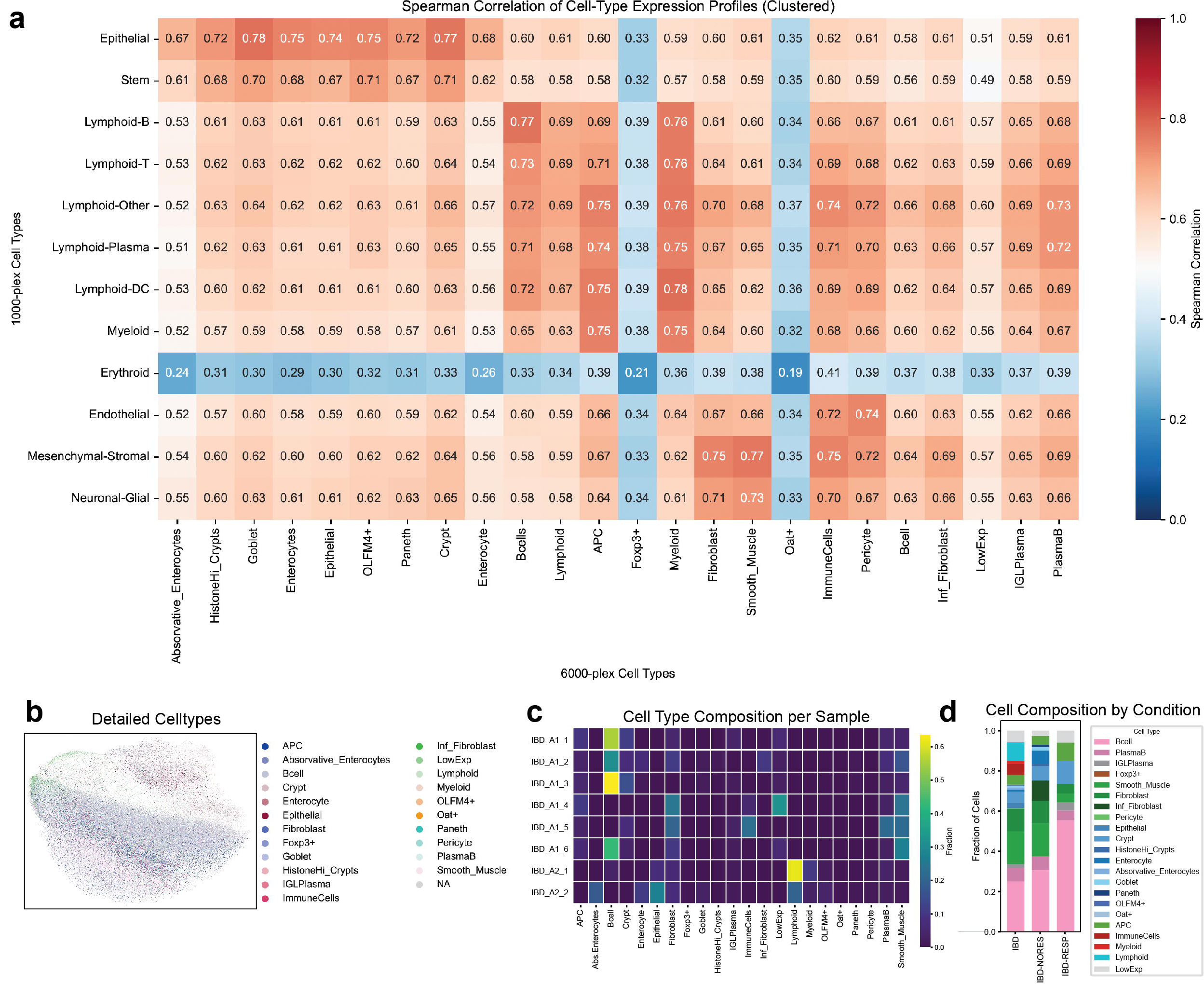

